# Regeneration-specific promoter switching facilitates Mest expression in the mouse digit tip to modulate neutrophil response

**DOI:** 10.1101/2024.06.12.598713

**Authors:** Vivian Jou, Sophia M. Peña, Jessica A. Lehoczky

## Abstract

The mouse digit tip regenerates following amputation, a process mediated by a cellularly heterogeneous blastema. We previously found the gene Mest to be highly expressed in mesenchymal cells of the blastema and a strong candidate pro-regenerative gene. We now show Mest digit expression is regeneration-specific and not upregulated in post-amputation fibrosing proximal digits. Mest homozygous knockout mice exhibit delayed bone regeneration though no phenotype is found in paternal knockout mice, inconsistent with the defined maternal genomic imprinting of Mest. We demonstrate that promoter switching, not loss of imprinting, regulates biallelic Mest expression in the blastema and does not occur during embryogenesis, indicating a regeneration-specific mechanism. Requirement for Mest expression is tied to modulating neutrophil response, as revealed by scRNAseq and FACS comparing wildtype and knockout blastemas. Collectively, the imprinted gene Mest is required for proper digit tip regeneration and its blastema expression is facilitated by promoter switching for biallelic expression.

## INTRODUCTION

Humans and mice can regenerate the distal region of the digits after amputation^1, 2^. This process involves the formation of a unique structure known as the blastema, which is the cellular source for the regenerated tissue and is a hallmark of epimorphic regeneration. The blastema is composed of numerous cell types, the majority of which are a heterogeneous population of fibroblasts, as we define by co-expression of markers Prx1, Pdgfrα, and Lum. Our prior study showed that while closely related by transcriptomics, the fibroblast population was composed of distinct subtypes suggesting separate biological functions^3^. We previously identified 67 genes enriched in the blastema fibroblasts^3^ and hypothesize that these are candidate pro-regenerative factors. Of these, we have prioritized Mest (mesoderm specific transcript), a member of the alpha/beta hydrolase family with structural similarity to epoxide hydrolases^4, 5^, though its substrate(s) is not defined.

Mest is a maternally imprinted gene first identified as a marker of mouse embryonic mesoderm and was originally named Peg1 (paternally expressed gene 1)^6, 7^. Beyond embryogenesis, Mest has been most studied in the context of adipogenesis and fat mass expansion in mouse obesity models^6, 8, 9^. Roles for Mest in tissue regeneration have recently been reported: Mest expression is associated with de novo adipogenesis in regenerative wound healing^10^, Mest is necessary for skeletal muscle regeneration following CTX-induced injury^11^, and expression of Mest can induce multipotency and increase differentiation potential in several cell types^12–14^. Corroborating these results, our blastema single-cell RNA sequencing (scRNAseq) data show Mest is expressed in fibroblasts with no defined computational lineage trajectory^15^. This suggests that Mest may function in defining early regenerative cell states during digit tip regeneration.

As has been found in both mouse and human, the Mest genomic locus is imprinted via methylation of the maternal allele, resulting in exclusive expression of the paternal allele^16, 17^. Monoallelic expression resulting from this imprinting is well-described for embryonic Mest expression and persists in adult tissues including blood, intestine, and adipose tissue^7, 18^. Biallelic expression of Mest has been reported in human blood lymphocytes and sporadically in the mouse spleen^7, 19^. However, for the Mest locus, this phenomenon is better studied in human cancers, specifically in breast, colorectal, and lung carcinomas and cell lines^20–23^. While Mest is strictly imprinted in embryos and often has biallelic expression in cancers, it is unknown how it is regulated during regeneration.

In this paper, we investigate the role that Mest, a maternally imprinted gene upregulated in the digit blastema, plays in mouse digit tip regeneration. We find that in vitro overexpression of Mest promotes bone differentiation, and in vivo Mest genetic loss of function results in delayed digit tip bone regeneration. Interestingly, paternal heterozygous knockout mice do not phenocopy the Mest homozygous knockout bone regeneration phenotype, but we establish that methylation of the imprinted region persists in the blastema. Instead, we find maternal allele expression arises via promoter switching to produce an alternate Mest transcript in regeneration, which does not occur during embryogenesis. Furthermore, through scRNAseq and FACs analyses, we find that neutrophil recruitment and clearance in the wounded tissue is impaired in the homozygous knockout mice, supporting a role for Mest in the inflammatory response. Ultimately, our data demonstrate that Mest is genetically necessary for proper digit tip regeneration and blastema Mest expression is facilitated by biallelic expression, despite genomic imprinting.

## RESULTS

### Increased Mest expression is regeneration-specific in the mouse digit

Our previous digit tip blastema single-cell RNA sequencing (scRNAseq) study identified Mest as a putative pro-regenerative factor^3^. We have now generated a 28 day post-amputation (dpa) digit tip scRNAseq sample (Figure S1A, B), the stage where regeneration is complete, and integrated the data with our original unamputated (UA), 11, 12, 14, and 17dpa scRNAseq dataset (Figure S1C-E). We computationally isolated and re-clustered the fibroblasts based on expression of Pdgfrα and Lum (Figures S1D and 1A), and Mest expression was found in a subset of the fibroblast subpopulations (Figure 1A’). Mest is highly upregulated in the blastemal fibroblasts from 11 to 17dpa compared to the unamputated and regenerated (28dpa) fibroblasts (Figure 1B).

**Figure 1.**
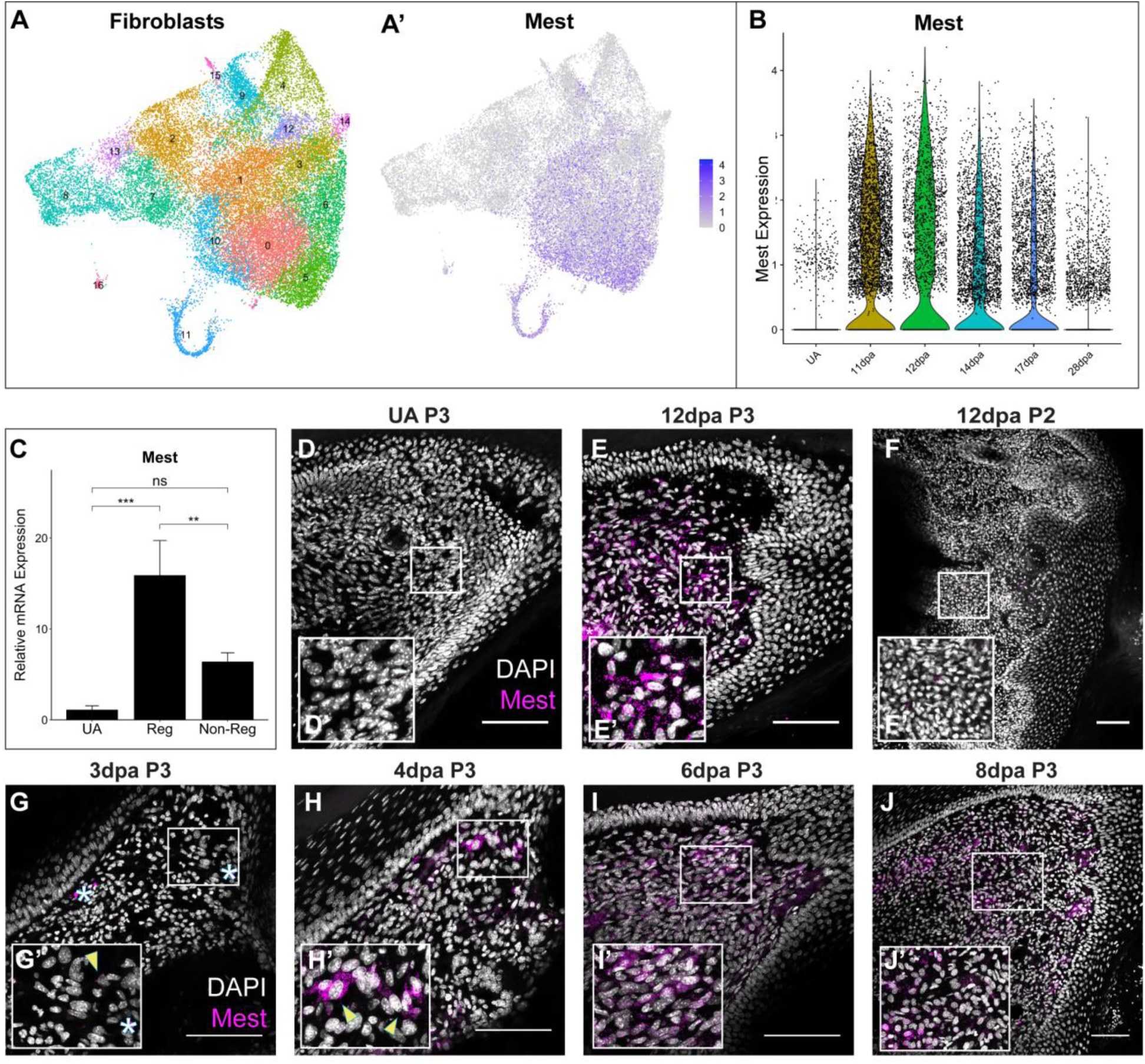
Mest is a candidate pro-regenerative factor. (A) UMAP plots of all-stages-integrated fibroblasts from the P3 digit showing Seurat clustering (A) and Mest expression (A’). (B) Violin plot of Mest expression in unamputated (UA), 11, 12, 14, 17, and 28dpa digit tip fibroblasts by scRNAseq. (C) qPCR analysis of Mest expression in UA distal digit, 12dpa regenerating P3 digit, and 12dpa non-regenerating P2 digit. Error bars depict standard deviation; (**) p < 0.01, (***) p < 0.001, ns = not significant. (D-J) Mest RNA expression (magenta) counterstained with DAPI (grayscale) HCR-FISH stains for UA (D) and regenerating P3 digit tip (E, G-J) and non-regenerative P2 proximal digit (F). White boxes show locations of inset panels (D’ - J’); all scale bars = 100µm. Yellow arrowheads denote example Mest expression; white asterisks denote autofluorescence.

To assess whether Mest upregulation is specific to blastema-mediated regeneration or more broadly in response to wound healing, we evaluated Mest expression in regenerative distal post-amputation digits (terminal phalanx; P3) compared to non-regenerative proximal post-amputation digits (second phalanx; P2) which undergo fibrotic wound healing^24, 25^. By qPCR analysis, Mest expression is increased 15X in the 12dpa blastema as compared to the UA digit mesenchyme (p=7.0e-4; Figure 1C). No statistically significant increase in Mest expression was found in 12dpa P2 fibrosing tissue as compared to the UA digit (p=0.095), and P2 expression was 2.5X less than P3 blastemal expression (p=0.007). These findings were corroborated by hybridization chain reaction RNA fluorescence in-situ hybridization (HCR-FISH)^26^; no appreciable Mest expression was detected in the P3 UA digit or the 12dpa P2 fibrosing digit while significant Mest expression was found in the blastema^3^ (Figure 1D-F, Figure S2A).

To determine the timing of Mest expression in the regenerating digit and if it is expressed earlier in the P2 non-regenerating tissue, we performed HCR-FISH at additional timepoints throughout regeneration and fibrosis. In the P3 digit tip, Mest is first expressed in a few cells adjacent to the amputation site between 3 and 4dpa (arrowheads, Figure 1G and H). The expression domain broadens by 6dpa, continues to increase through the blastema stages, then decreases as the blastema matures and regeneration is completed (Figure 1E, G-J, Figure S2A-C). In contrast, no Mest expression was found by HCR-FISH during early wound healing in the P2 non-regenerating digit at 3, 6, or 9dpa (Figure S2D-F), correlating upregulation of Mest with blastema-mediated regeneration, not wound healing.

### Mest does not induce multipotency for mesenchymal stem cell lineages in vitro

We performed computational lineage trajectory analysis with the integrated and subsetted fibroblast scRNAseq data using SPRING^15^ (Figure 2A). From this, we can see that the fibroblasts in the UA digit tip project away from the blastema stage cells, indicating significant transcriptomic differences between homeostatic and regenerative fibroblasts. Additionally, the newly added 28dpa fibroblasts project similar to the UA fibroblasts, supporting a return to a homeostatic state (Figure 2A’). Several differentiation trajectories are predicted, including osteoblastic and other musculoskeletal and connective tissue lineages (Figures 2A’’ and S2G-J), consistent with our previous report^3^. Mest expression is associated with cells of early blastema stages, not the differentiating cells (Figure 2A’’) suggesting Mest expression is upregulated in undifferentiated and progenitor-like cells.

**Figure 2.**
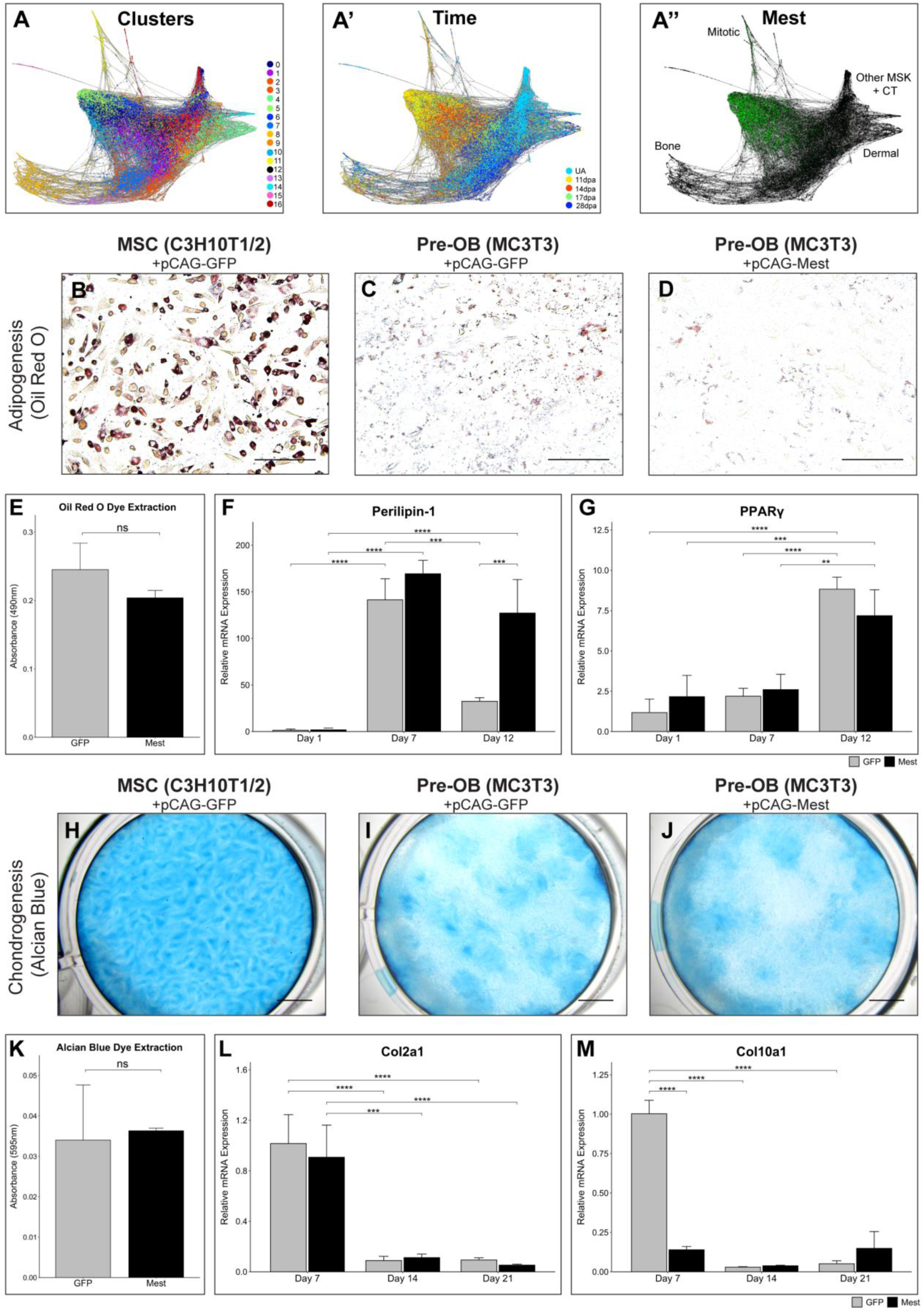
No Mest-induced in vitro MSC multipotency. (A) SPRING plot of the integrated scRNAseq fibroblasts, (A) colored by Seurat clusters, (A’) time point, and (A’’) Mest expression in green; MSK = musculoskeletal; CT = connective tissue. (B-D) Oil Red O staining of (B) pCAG-GFP C3H10T1/2, (C) pCAG-GFP MC3T3, and (D) pCAG-Mest MC3T3 cells at 12 dpi. (E) Quantification of extracted Oil Red O from 12dpi stained MC3T3 cells in (C) and (D). (F-G) Adipogenesis qPCR analysis of (F) Perilipin-1 and (G) PPARγ genes in GFP (gray) and Mest (black) transfected MC3T3 cells at 1, 7, and 12dpi. (H-J) Alcian Blue staining of (H) pCAG-GFP C3H10T1/2, (I) pCAG-GFP MC3T3, and (J) pCAG-Mest MC3T3 cells at 21dpi. (K) Quantification of Alcian Blue stain extracted from 21dpi stained MC3T3 cells in (I) and (J). (L-M) Chondrogenesis qPCR analysis of (L) Col2a1 and (M) Col10a1 genes in GFP and Mest transfected MC3T3 cells at 7, 14, and 21dpi. Error bars depict standard deviation. Statistical comparisons are shown for those discussed in text; (**) p < 0.01, (***), p < 0.001, (****) p < 0.0001, ns = not significant. Scale bars in images (B-D) = 100µm, scale bars in images (H-J) = 250µm.

Recent reports find in vitro overexpression of Mest in mouse adipocytes or human periodontal ligament cells induces transdifferentiation^13, 14^. Thus, we hypothesized that Mest expression in digit tip fibroblasts could induce multipotency for fibroblastic lineages. To test this, we utilized immortalized pre-osteoblasts (MC3T3) to determine if Mest overexpression could induce multipotency for adipogenesis and chondrogenesis, mesenchymal stem cell differentiation lineages distinct from osteogenesis (Figure S3A). MC3T3 cells were transfected with a constitutively expressing Mest construct (pCAG-Mest) or a control GFP construct (pCAG-GFP) and were compared with immortalized mouse mesenchymal stem cells (MSCs; C3H10T1/2) transfected with pCAG-GFP as a multipotency control. Mest RNA and protein expression was verified 24-48 hours post Mest transfection (Figure S3B-D). Oil Red O staining for terminal adipogenesis 12 days post-induction (dpi) indicated robust differentiation of the control GFP transfected C3H10T1/2 MSCs and no significant differentiation in the control GFP MC3T3 pre-osteoblasts (Figure 2B, C). Similarly, no significant terminal adipogenesis occurred in the context of Mest overexpression (Figure 2D, E). To determine if Mest overexpression induced early adipocyte identity, 1, 7, and 12dpi RNA was collected and evaluated for adipogenesis marker genes Perilipin-1 and PPARγ^27^. Perilipin-1 expression in control C3H10T1/2 cells at 7dpi increased 2,000X (p=2.4e-5) and significantly decreased by 12dpi (p=3.3e-5; Figure S3E); PPARγ expression increased by 30X at 12dpi (p=0.001) (Figure 2F, G). While both GFP and Mest transfected MC3T3 cells expressed Perilipin-1 by 7dpi, this increase was significantly less than the 2000X increase seen in C3H10T1/2 cells. Interestingly, while Perilipin-1 expression decreased by 12dpi in GFP MC3T3 cells, it was maintained in Mest transfected cells, suggesting persistence of an early adipogenic-like state (Figure 2F). The expression of PPARγ was similar in both GFP and Mest expressing MC3T3 cells with only a 7.5X increase in gene expression at 12dpi (Figure 2G). The absence of a significant difference in PPARγ expression between the Mest and GFP overexpressing cells at 12dpi suggests the cells express low levels of adipogenic genes in response to the differentiation media, but do not truly differentiate.

To assess if Mest overexpression in pre-osteoblasts can generate progenitors competent for chondrogenesis we assessed terminal differentiation by Alcian Blue staining at 21dpi. Robust chondrogenesis of the control C3H10T1/2 cells was found (Figure 2H) though minimal differentiation of either GFP or Mest transfected MC3T3 cells occurred (Figure 2I-K). RNA isolated at 7, 14, and 21dpi was assessed for early and late chondrogenesis marker genes Col2a1 and Col10a1^28^. Control C3H10T1/2 cells had peak expression of Col2a1 at 7dpi that decreased 3X by 21dpi while Col10a1 expression exhibited a 7.5X increase at 14dpi (p=0.02) and persisted through until 21dpi (Figure S3G, H). Both GFP and Mest overexpressing MC3T3 cells had a peak expression of Col2a1 at 7dpi with a sharp decline in expression at 14 and 21dpi (Figure 2L). However, neither had increased Col10a1expression by 21dpi, supporting the negligible chondrogenesis found by Alcian Blue staining (Figure 2M, S3H). These data demonstrate that overexpression of Mest in pre-osteoblasts does not induce a mesenchymal stem cell-like multipotency competent for adipogenesis or chondrogenesis in vitro.

### Mest expression modulates osteogenesis in vitro and in vivo

While Mest over-expression did not induce multipotency of pre-osteoblasts for other MSC lineages, we found it changed the timing and amount of osteogenesis in vitro. 28dpi cells stained with Alizarin Red revealed a 6.8X increase in ossification in the Mest versus GFP transfected C3H10T1/2 cells (p=0.004; Figure 3A-C), which was also observed for MC3T3 cells (Figure S3I, J). Consistent with these findings, Runx2 expression at 7dpi was 2.5X higher in Mest expressing cells as compared to GFP (p=0.04; Figure 3D), demonstrating that Mest promotes osteogenesis in both mouse pre-osteoblasts and MSCs.

**Figure 3.**
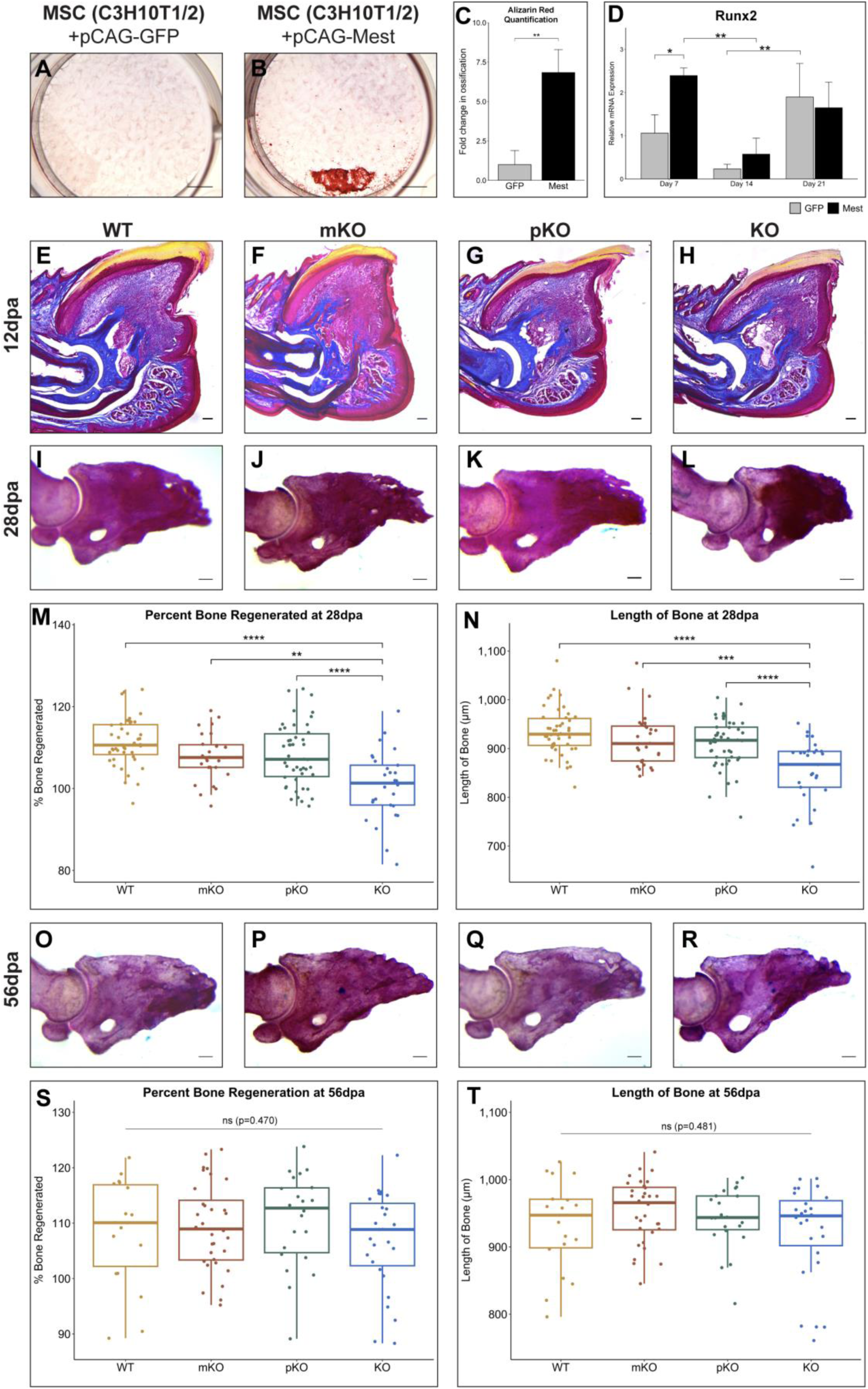
Mest promotes osteogenesis. (A-B) Alizarin Red staining of (A) pCAG-GFP and (B) pCAG-Mest transfected C3H10T1/2 cells at 28dpi. (C) Quantification of Alizarin Red stained C3H10T1/2 cells at 28dpi in Fiji. (D) qPCR analysis of osteogenic marker Runx2 in GFP (gray) and Mest (black) transfected cells at 7, 14, and 21dpi. Statistical comparisons are shown for those discussed in text. (E-H) Representative 12dpa Masson trichrome stained digit sections of (E) Mest-WT, (F) mKO, (G) pKO, and (H) KO mice. (I-L) Representative whole-mount Alizarin Red staining of (I) Mest-WT, (J) mKO, (K) pKO, and (L) KO 28dpa digits. (M-N) Box plots with individual data points shown of the percent bone regenerated (M) and length of P3 bone (N) for Mest-WT, mKO, pKO, and KO 28dpa Alizarin Red stained digits. Percent bone regeneration was calculated as area of each P3 digit/average area of unamputated digits for that respective genotype. (O-R) Representative whole-mount Alizarin Red staining of (O) Mest-WT, (P) mKO, (Q) pKO, and (R) KO 56dpa digits. (S-T) Box plots with individual data points shown of (S) the percent bone regenerated and (T) length of P3 bone for Mest-WT, mKO, pKO, and KO 56dpa Alizarin Red stained digits. Error bars depict standard deviation; (*) p < 0.05, (**) p < 0.01, (***) p < 0.001, (****) p < 0.0001, ns = not significant. Scale bars in images (A) and (B) = 250µm, all other scale bars = 100µm.

The ability of Mest to modulate in vitro osteogenesis suggests the expression of Mest in the blastema may regulate digit tip bone regeneration. To determine if genetic deletion of Mest yields a digit tip regeneration phenotype, we utilized a characterized Mest knockout allele^29^. Mest is a maternally imprinted gene^4, 16^, thus we analyzed digit tip regeneration in the following genotypic cohorts: Mest wildtype (WT), heterozygous maternal knockout (mKO), heterozygous paternal knockout (pKO), and homozygous knockout (KO). Weights of 8-10 week-old mice did not differ between genotype cohorts within each sex (Figure S4A, B) supporting no gross developmental differences. Masson trichrome histology of 12dpa digits revealed that all Mest genotypes formed blastemas that did not statistically differ in 2D area when averaged across at least 6 blastemas per genotype (Figure 3E-H, S4C). Masson trichrome histology at 28dpa revealed that all tissues regenerated for all genotypes, with digits consisting of a full-length nail, bone, epithelium, and mesenchymal tissue surrounding the bone (Figure S4D-G). Additional cohorts were analyzed by whole mount Alizarin Red staining at 28dpa, for which we measured the 2D area and length of each P3 bone (Figure 3I-L, Figure S5A-D). While the unamputated digits across genotypes did not significantly differ in either measurement (Figure S5E, F), 28dpa Mest-KO regenerated digit tip bones were significantly shorter (WT vs KO p=3.6e-9) and exhibited less total bone regeneration (WT vs KO p=2.3e-8) as compared to all other genotypes (Figure 3M, N, S5G). Interestingly, digits analyzed at 56dpa, a stage well-beyond the completion of regeneration, revealed that Mest-KO digits had caught up with the other genotypes as assessed by percent bone regeneration and length (Figure 3O-T). This time point also revealed that the unamputated 56-day Mest-KO bones were significantly smaller and shorter than the other genotypes, suggesting a potential defect in attaining peak bone mass and length between 12-16 weeks (Figure S5H-M). Importantly, this phenotype is separable from regeneration; while the 2D bone area of the 56dpa Mest-KO regenerated bones was significantly smaller than the other genotypes (Figure S5N), the digits regenerate normally as compared to the UA controls (Figure 3S, T). Taken together, these genetic experiments indicate that Mest is not necessary for blastema formation, but functions in blastema osteogenic differentiation, corroborating our in vitro findings.

### Genomic imprinting of the Mest promoter persists in the digit tip blastema

Previous studies have established that the Mest genomic locus is maternally imprinted via methylation around the first exon during embryonic development as well as in adult Mest expressing tissues^7, 16^ (Figure 4A). Biallelic expression of Mest has been reported in primary human breast and colorectal cancers and lung cancers, both primary and cell lines, and at low frequencies in another mouse Mest knockout allele^20–23, 30, 31^. Within this framework, we anticipated the Mest-pKO digits to phenocopy the Mest-KO cohort; however, no significant difference was found in percent regeneration or bone length at 28dpa among the WT, mKO, and pKO groups (Figure 3M, N). We first verified this was not secondary to expression changes in miR-335, a microRNA embedded in intron 2 of the Mest locus shown to promote osteogenesis^32, 33^ (Figure 4A, B). With no significant difference in miR-335 expression among genotypes (ANOVA p=0.122), the absence of Mest-pKO bone regeneration phenotype instead suggests continued expression of Mest in the paternal knockout regenerating digits. qPCR analysis confirms that there is no significant difference in WT and mKO 12dpa Mest expression, and that KO blastema expression is 16.7X lower than WT (p=8.4e-5; Figure 4C). The pKO blastema expression is 3X less than that of WT blastemas (p=8.7e-4) and 5.7X higher than KO, though not statistically significant (p=0.19; Figure 4C), which does not align with expectations for an imprinted gene. Moreover, the qPCR primers are specific to sequences within the knockout allele deletion (Figure 4A); thus, any amplification is de facto from the wildtype allele. Therefore, Mest expression in the Mest-pKO 12dpa blastema is derived from the maternal allele, supporting biallelic expression of Mest during digit tip regeneration.

**Figure 4.**
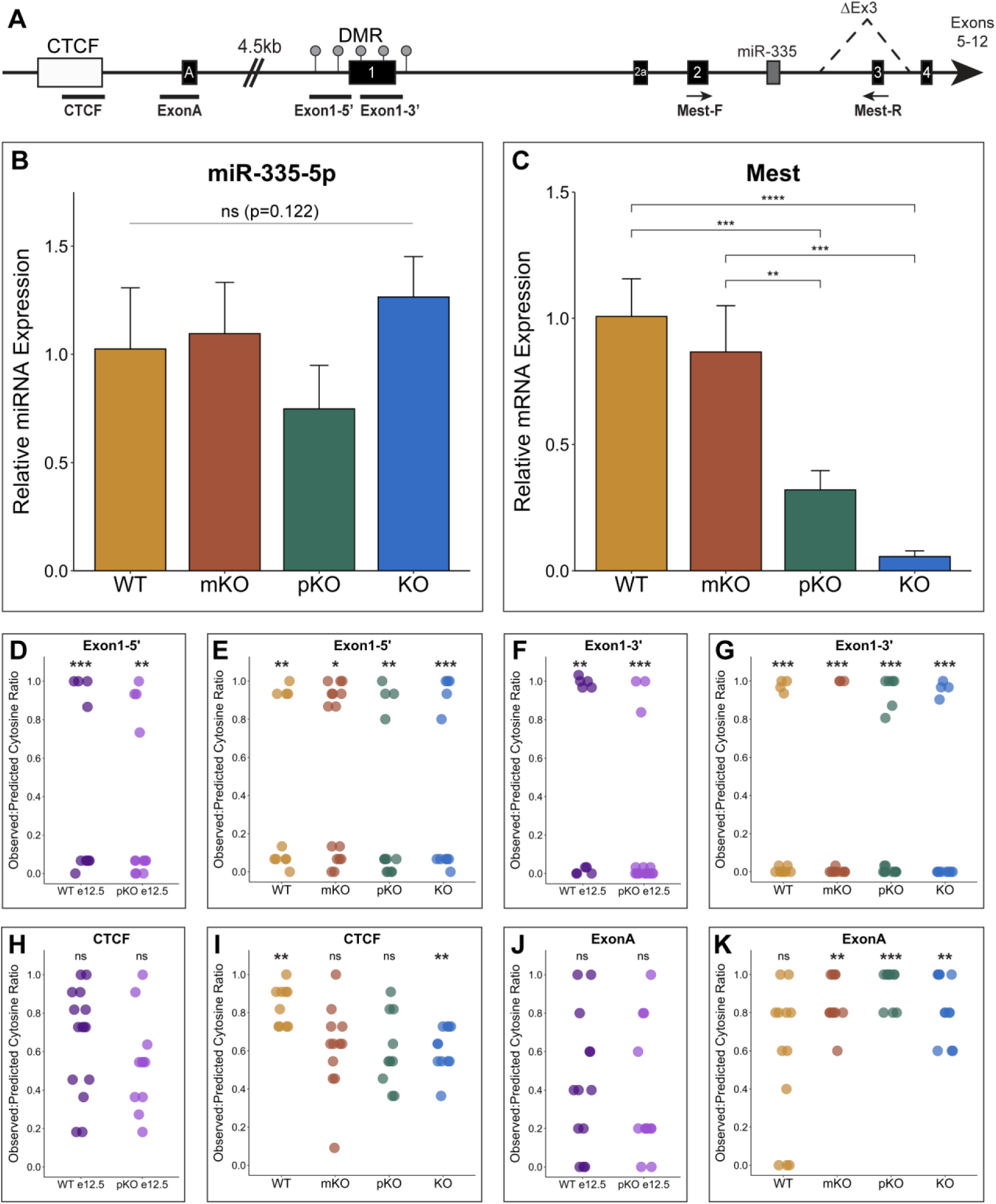
Genomic imprinting of Mest persists in the blastema. (A) Schematic of the Mest genomic locus with exons as black boxes with exon number labeled in white; ΔEx3 denotes Mest-KO deletion. DMR = differentially methylated region (gray circles); CTCF = CTCF element (white box); miR335 = microRNA-335 (dark gray box); black bars denote regions of bisulfite sequencing. Location of Mest qPCR primers shown with black arrows. (B) microRNA-335-5p qPCR analysis in Mest-WT, mKO, pKO, and KO 12dpa blastemas. (C) Mest qPCR in Mest-WT, mKO, pKO, and KO 12dpa blastemas. (D-K) Dot plots of bisulfite sequencing observed:predicted cytosine ratios for CTCF, ExonA, and Exon1 PCR amplicons shown in (A); each dot represents ratio for one sequenced clone. A ratio of 1 is fully methylated and a ratio of 0 is not methylated. (D, F, H, J) Bisulfite sequencing clones from embryonic e12.5 hindlimb Mest-WT and pKO DNA. (E, G, I, K) Bisulfite sequencing clones from 12dpa blastemas of Mest-WT, mKO, pKO and KO mice. (*) p < 0.05, (**) p < 0.01, (***) p < 0.001, (****) p < 0.0001, ns=not significant.

To explore potential loss of imprinting of the maternal allele in digit tip regeneration, we used targeted bisulfite sequencing to assess methylation surrounding the canonical promoter (Figure 4A). A differentially methylated region (DMR) surrounds exon 1, also referred to as a CpG island, due to hypermethylation of the maternal allele and hypomethylation of the paternal allele (Figure 4A)^16^. We assayed the upstream and downstream regions of this DMR, Exon1-5’ and Exon1-3’ (Figure 4A), using bisulfite converted genomic DNA isolated from Mest-WT, mKO, pKO, and KO 12dpa digit tip blastemas, as well as from Mest-WT and pKO e12.5 embryonic hindlimbs as controls. Cloned PCR amplicons were sequenced and scored for observed to predicted methylated cytosine ratios, whereby a ratio nearing 0 reflects hypomethylation and a ratio nearing 1 reflects hypermethylation. Consistent with imprinting of the Mest locus during development, cytosine ratios for Exon1-5’ and Exon1-3’ clones from e12.5 hindlimb DNA were either hypo- or hypermethylated, even in the context of the paternal knockout (Figure 4D, F). Importantly, this bimodal methylation of alleles was also found for Mest-WT 12dpa digit tip blastemas, demonstrating that genomic imprinting of Mest continues during digit tip regeneration (Figure 4E, G). Moreover, Mest genetic loss of function does not change methylation of the canonical promotor as seen by Mest-mKO, pKO, and KO 12dpa blastemas (Figure 4E, G S6A, B). These data demonstrate persistent genomic imprinting of Mest when it is highly expressed during regeneration, even under the pressure of genetic loss of function.

With genomic imprinting intact, we reasoned that biallelic Mest expression found in the pKO blastema could be regulated by an alternate promoter. Indeed, an alternate transcription start site exists 4.5kb upstream of the Mest canonical promoter (Figure 4A)^18, 34^. We focused on two regions to characterize DNA methylation: one within a CTCF binding element conserved with the human locus and one covering exon A, where another CpG island exists in humans but not mice^35^. We evaluated methylation of these two regions, CTCF and ExonA (Figure 4A, S6C), by bisulfite sequencing. Clones through both regions in Mest-WT and pKO e12.5 hindlimbs revealed a uniform distribution of allelic methylation patterns, indicating that this upstream promoter is not imprinted or strictly regulated by methylation during limb development (Figure 4H, J). Methylation of ExonA in the Mest-WT 12dpa blastema was similarly uniform, and all amplicons through the CTCF element were significantly hypermethylated (Figure 4I, K). However, unlike the embryonic limb, methylation of the upstream Mest promoter in the 12dpa blastema changes in the context of Mest genetic loss of function. The CTCF region becomes increasingly hypomethylated in the Mest-mKO, pKO, and KO blastemas, and this demethylation is most dramatic in the KO blastema with an average cytosine ratio of 0.61 compared to 0.84 in the WT blastema (Figure 4I and S6D). The ExonA region becomes increasingly hypermethylated in Mest-mKO, pKO, and KO blastemas (Figure 4K), specifically in CpG positions directly upstream of the transcription start site for the Mest210 variant (Figure S6C, E). However, with only 5 predicted methylated CpGs for the ExonA amplicon, interpretation of this finding is challenging. Collectively, these data highlight a change in gene regulation at an upstream promoter as a mechanism to compensate for Mest genetic loss of function during regeneration, suggesting that Mest-pKO blastema biallelic expression may result from this change in upstream regulation.

### Biallelic Mest expression during regeneration occurs via promoter switching

To determine whether this shift in methylation of the Mest upstream promoter impacts transcription, we analyzed three Mest transcript variants: Mest202, Mest210, and Mest211 (NM_001252292, NM_001252293, and NM_008590). Mest202 and Mest211 are both regulated by the imprinted exon 1 promoter region and Mest210 is regulated by the alternate upstream exon A promoter region (Figures 4A, 5A)^16, 18^. These variants differ in transcription and translation start sites, though they all encode the same protein with minor variation in the N-terminus (Figure 5A and S7A). Importantly, the MEST N-terminus is not predicted to contain any functional domains and all three variants encode the downstream enzymatic domain, supporting conserved function among them (Figure S7B).

**Figure 5.**
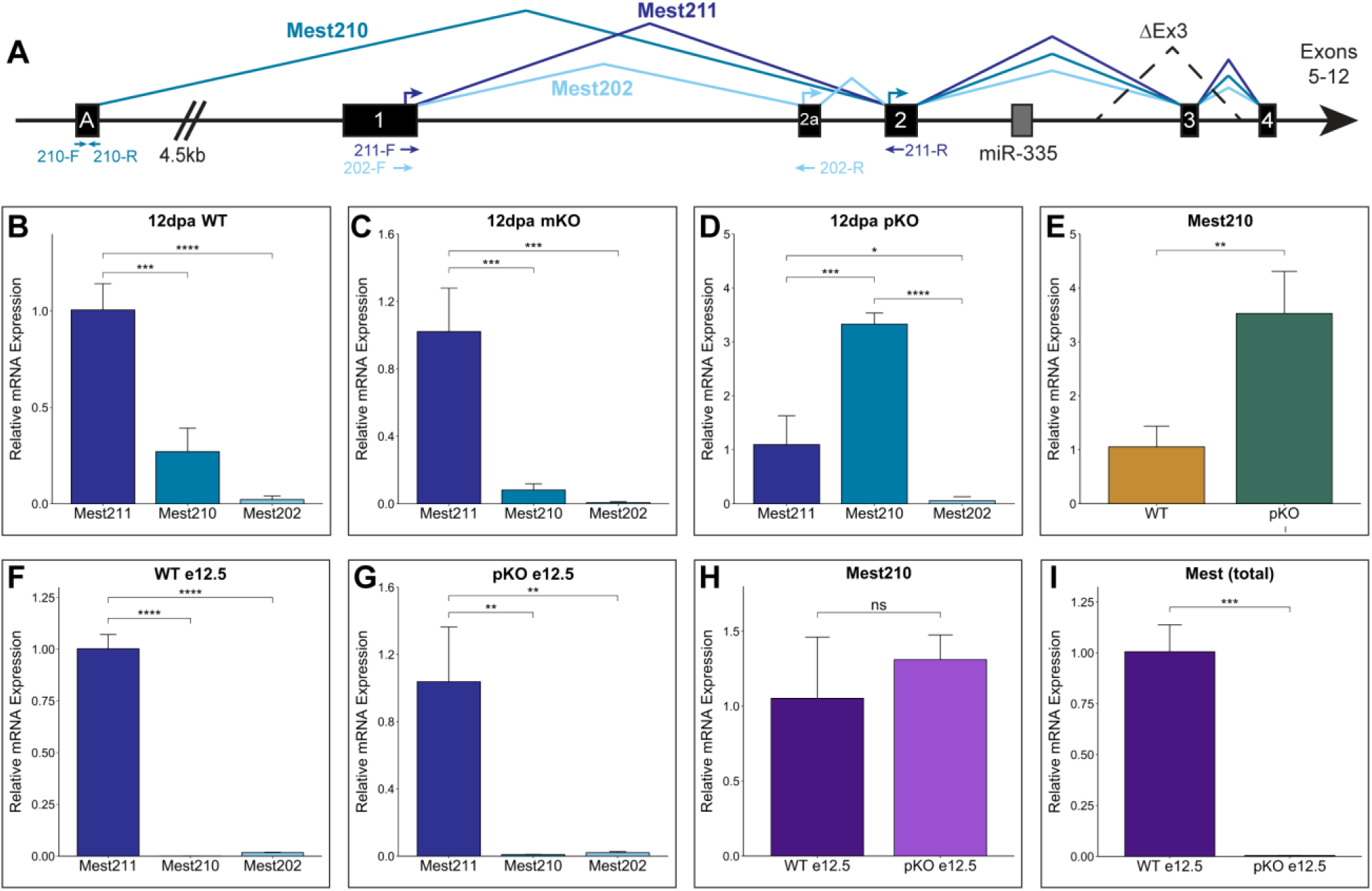
Biallelic expression of Mest via promoter switching. (A) Schematic of Mest locus with three annotated transcript variants: Mest211 (NM_008590; dark blue), Mest210 (NM_001252293; blue), and Mest202 (NM_001252292; light blue). Position of transcript-specific qPCR primers shown with arrows of their respective colors below the schematic and arrows depicting translation start sites are above the exons. (B-D) Relative expression levels of each variant by qPCR analysis in (B) Mest-WT, (C) mKO, and (D) pKO 12dpa blastemas. (E) qPCR analysis of Mest210 expression in Mest-WT and pKO 12dpa blastemas. (F, G) qPCR analysis of transcript variant expression in (F) Mest-WT and (G) pKO e12.5 embryos. (H) qPCR analysis of Mest210 expression in Mest-WT and pKO e12.5 embryos. (I) Total Mest expression in Mest-WT and pKO e12.5 embryos using primers shown in Figure 4A. Error bars are reported as standard deviation; (*) p < 0.05, (**) p < 0.01, (***) p < 0.001, (****) p < 0.0001.

Using variant-specific qPCR, we assessed Mest transcript variant expression in 12dpa blastemas isolated from the Mest genotypic cohorts. We found that in the wildtype blastema, Mest211 is the predominant isoform, with significantly less Mest210 and Mest202 expression (3.7X and 47X lower, respectively; Figure 5B). This variant expression profile was highly similar in Mest-mKO blastemas, which only express Mest from the paternal allele (Figure 5C). In contrast, Mest-pKO blastemas predominantly express the Mest210 variant, with a 3.3X increase compared to canonical Mest211 (p=5.3e-5; Figure 5D), and 3.5X more Mest210 than Mest-WT blastemas (p=0.008; Figure 5E). The increased Mest210 expression in Mest-pKO blastemas is consistent with maternal allele expression derived from exon A that undergoes epigenetic changes (Figure 5A, 4K), circumventing the imprinted Mest211 promoter (Figure 5A, 4D-G). These analyses also revealed that Mest202 is the minor Mest variant, with extremely low expression in the blastema for all genotypes (Figure 5B-D).

To determine if Mest promoter switching is unique to the blastema, we assessed these transcript variants in Mest-WT and pKO e12.5 embryos, when Mest is highly expressed and strictly imprinted. As found for the blastema, canonical Mest211 is the predominant Mest transcript in the wildtype embryos, with significantly less Mest210 expression (1000X lower) and Mest202 expression (59X lower) (Figure 5F). In stark contrast to the Mest-pKO 12dpa blastema, Mest210 transcription is not upregulated in pKO e12.5 embryos. Mest210 and Mest202 exhibit 105X and 48X lower expression relative to Mest211, respectively (Figure 5G), and there is no significant difference in Mest210 expression between WT and pKO e12.5 embryos (p=0.367, Figure 5H). These data align with the lack of difference in Exon1, ExonA, and CTCF methylation profiles between Mest-WT and pKO embryos (Figure 4D, F, H, J), ultimately resulting in negligible Mest expression in pKO embryos (Figure 5I). These findings support previous studies establishing strict imprinting during development^7, 19^ and further illustrate promoter switching to Mest210 is specific to regeneration.

### Mest knockout blastemas have impaired neutrophil response during inflammation

To begin to understand the function of Mest and the need for its high expression during digit tip regeneration, we performed single-cell RNA transcriptomics on Mest-WT and Mest-KO 12dpa blastemas. Post-quality control and filtering, our dataset included 8,802 wildtype cells and 6,898 knockout cells. All digit tip blastema cell types previously identified by scRNAseq^3, 36^ were present in both the WT and KO samples (Figure S8A). Integration of the WT and KO data revealed the two genotypes were well-mixed, confirming that Mest-KO indeed form a blastema (Figure 3H, S8A, B). All major cell types have proportional population sizes except for epithelial cells, though this is secondary to our dissection technique to minimize epithelium (Figure S8C). We subsetted and re-clustered the integrated fibroblasts for more detailed analysis since this cell type highly expresses Mest (Figure 6A, S8D). We looked for differentially expressed genes (DEGs) between the Mest-WT and KO fibroblasts with an average log2FC > 0.58 and expressed in at least 25% of the cells, and found 119 significantly upregulated genes in the Mest-WT fibroblasts and 22 upregulated genes in the Mest-KO fibroblasts (Figure 6B, Table S2, S3). Of the top 10 upregulated genes in the wildtype fibroblasts, four are chemokines involved in leukocyte recruitment, especially neutrophils (Ccl2, Cxcl5, Cxcl1, and Cxcl2)^37^, and other genes including Tnfaip3 and Nfkbia are involved in regulating inflammation^38, 39^ (Table S2). Gene set enrichment analysis of the 119 upregulated Mest-WT DEGs established the IL-17 and TNF signaling pathways were significantly upregulated, consistent with leukocyte recruitment (Figure 6C). To assess which fibroblast subpopulation(s) may be driving the inflammation associated DEGs, we used differential proportion analysis and found significantly fewer Mest-KO fibroblasts in cluster-0 as compared to Mest-WT (p=0.007; Figure 6D). Indeed, KEGG pathway analysis of cluster-0 marker genes filtered by average log2FC > 1.0 and expressed in at least 25% of cells revealed enrichment of IL-17 and TNF signaling among other pathways (Figure 6E), and the chemokine DEGs we previously identified (Figure 6B) are differentially expressed between Mest-WT and KO cluster-0 fibroblasts (Figure S8E, Table S4). These findings suggest that Mest-KO blastema fibroblasts have reduced neutrophil chemoattractant production, primarily due to a reduction in the cluster-0 subpopulation.

**Figure 6.**
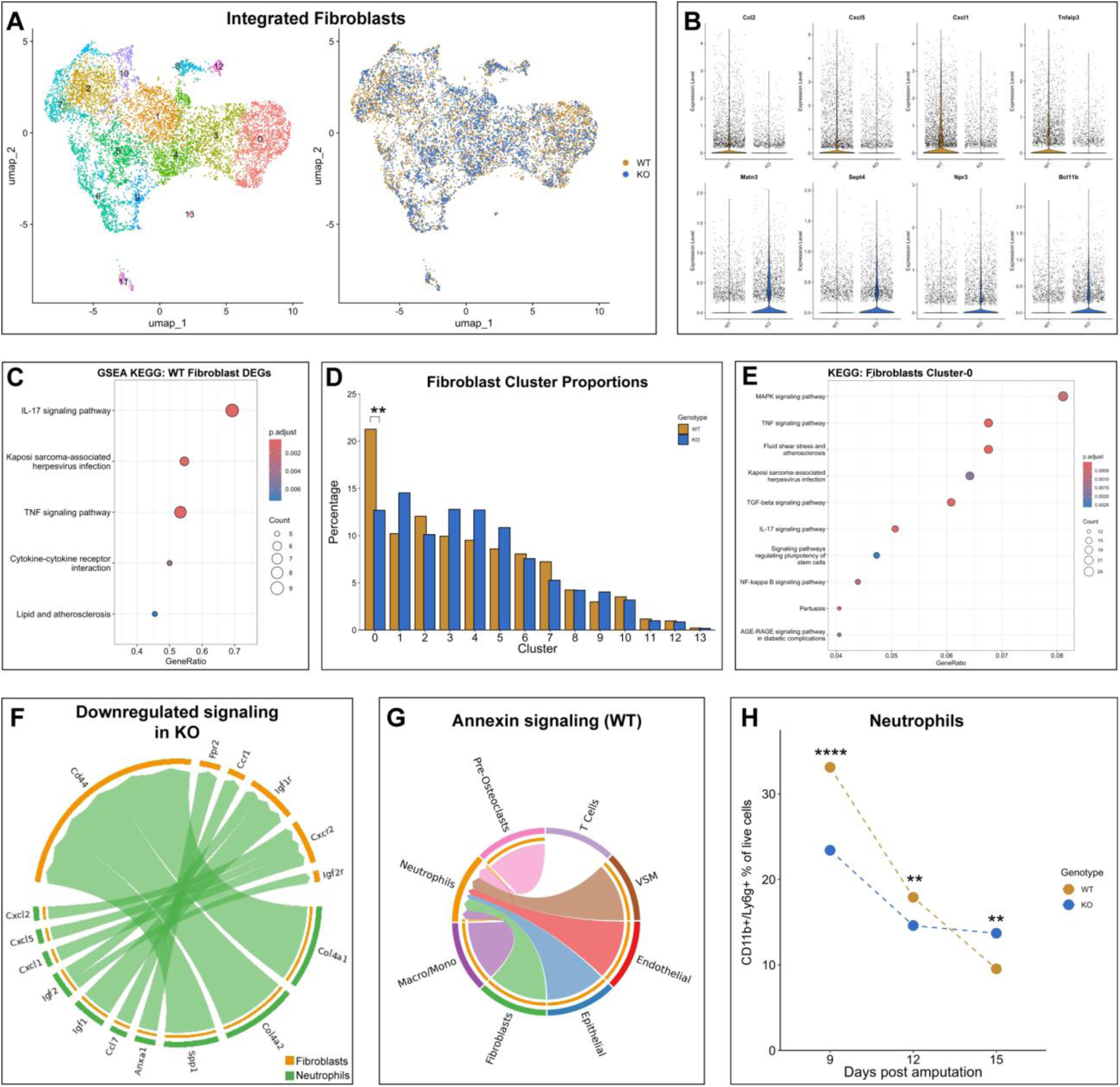
Single-cell transcriptomic analysis of Mest wildtype and knockout blastemas. (A) UMAP plot of Mest-WT and KO 12dpa integrated blastema fibroblasts colored by Seurat clusters (left) and by genotype (right). (B) Violin plots of top 4 differentially expressed genes (DEGs) identified between WT (top) and KO fibroblasts (bottom). (C) Dot plot of gene set enrichment analysis using KEGG pathways derived from Mest-WT DEGs. (D) Relative population proportions for each genotype in the integrated fibroblast dataset. (E) Dot plot of KEGG analysis derived from cluster-0 DEGs. (F) Chord plot for ligand-receptor pairs between fibroblasts and neutrophils that are downregulated in the Mest-KO blastema compared to WT. (G) Chord plot for Annexin signaling by cell type in the WT blastema. (H) Mest-WT and KO blastema neutrophils quantified by flow cytometry at 9, 12, and 15dpa. Percentages were calculated for CD11b^+^/Ly6g^+^ cells out of all PI^-^ cells. (**) p < 0.01, (****) p < 0.0001.

Finding differential inflammatory signaling between Mest-WT and KO fibroblasts led us to characterize the immune cells at a higher resolution. We subsetted and re-clustered the macrophages and monocytes, neutrophils, T cells, and pre-osteoclasts and found that the largest subpopulation change was in the neutrophils, in which the Mest-KO blastema had fewer neutrophils than the Mest-WT blastema (Figure S8F, G; cluster 2 red arrow). This reduction was not statistically significant by differential proportion analysis (p=0.117) though it conceptually aligns with our finding of reduced expression of neutrophil recruitment chemokines in the Mest-KO fibroblasts. We used cell-cell communication computational tools CellChat^40^ and NICHES^41^ to identify signaling relationships between fibroblasts and neutrophils. Both tools corroborated the decreased cytokine interactions between fibroblasts and neutrophils in the Mest-KO blastema (Figure 6F and S9A, B). In addition, we found that Annexin (Anxa1) signaling, a key pathway involved in the resolution of inflammation^42^, occurs in the wildtype blastema between neutrophils and many cell types including fibroblasts (Figure 6G). However, Anxa1 signaling is absent in the knockout blastema, further underscoring a disrupted immune response in Mest-KO digits (Figure 6G and S9A-D, red arrows).

To evaluate these findings in vivo, we performed flow cytometry for neutrophils in Mest-WT and KO blastemas at 9, 12, and 15dpa (Figure 6H, S10A, B, Table 1). Consistent with our scRNAseq finding of reduced production of leukocyte chemoattractants, there were significantly fewer CD11b+/Ly6g+ neutrophils at 9 and 12dpa in the Mest-KO blastemas as compared to Mest-WT (p=0.000 and p=0.005, respectively; Figure 6H). However, by 15dpa there were significantly more CD11b+/Ly6g+ neutrophils in the Mest-KO blastemas as compared to Mest-WT (p=0.003), suggesting neutrophil clearance as required for the resolution of inflammation is delayed^43^. Taken together, these data indicate that the Mest-KO blastema has an impaired immune response due to reduced chemokine production in a subpopulation of fibroblasts, which prolongs the inflammatory process and ultimately delays digit bone regeneration.

**Table 1.** Neutrophil flow cytometry results. Raw cell numbers for all PI^-^ cells and double positive CD11b^+^/Lyg6^+^ cells from flow cytometry analysis for WT and KO tissues at 9, 12, and 15dpa.

|  |  | All PI<sup>-</sup> cells | CD11b<sup>+</sup>/Lyg6<sup>+</sup> cells | p-value |
| --- | --- | --- | --- | --- |
| **9dpa** | WT | 18275 | 6044 | 0 |
|  | KO | 20476 | 4785 |  |
| **12dpa** | WT | 31867 | 5697 | 0.00447 |
|  | KO | 32874 | 4799 |  |
| **15dpa** | WT | 17387 | 1659 | 0.00257 |
|  | KO | 18465 | 2530 |  |

## DISCUSSION

We began this study with the hypothesis that Mest is a pro-regenerative gene that can drive plasticity of fibroblast lineages during digit tip regeneration. However, our in vitro experiments show that Mest overexpression in pre-osteoblasts is not sufficient to induce a multipotent progenitor state capable of differentiating into adipogenic or chondrogenic lineages (Figure 2). This data conflicts with other reports where Mest did induce lineage transdifferentiation, though the differences may be due to distinct species, cell types, or lineages assayed^13, 14^. Interestingly, the control in vitro osteogenic differentiation assays in our experiments revealed Mest overexpression significantly increased osteogenesis (Figure 3A-D, S3I, J), which ultimately complemented the delayed digit tip bone regeneration we found in the Mest-KO mice (Figure 3I-T). This finding is nuanced because it is the composite of two distinct bone phenotypes: failure to attain peak bone mass/length and reduced bone regeneration. At 28dpa (12 weeks old), Mest-KO digit tips have reduced bone regeneration, but the UA digits support normal bone development among genotypes. However, by 56dpa (16 weeks old), the Mest-KO UA digits are significantly smaller in area and length than the other genotypes, suggesting a failure to reach peak bone mass/length which occurs around this time on the C57Bl6/J background^44, 45^. Within the context of a smaller digit tip bone, the Mest-KO bones have fully regenerated by 56dpa, consistent with delayed regeneration.

The mechanism of how Mest expression regulates osteogenesis and bone regeneration remains to be determined, though our scRNAseq experiments suggest this is an indirect effect. We found that Mest expression in a subpopulation of fibroblasts mediates cytokine production and ultimately neutrophil recruitment and clearance following digit tip amputation (Figure 6, Table 1), likely resulting in an environment of delayed regeneration. Broadly speaking, the resolution of inflammation, which includes clearance of neutrophils, is a critical phase of wound-healing and tissue repair^43, 46^. Our scRNAseq cell-cell communication analyses highlight Annexin signaling, a key pathway in initiating the resolution of inflammation^42, 46^, is absent in Mest-KO blastemas (Figure 6G, S9A), which aligns with more neutrophils in Mest-KO 15dpa blastemas (Figure 6H). Intriguingly, MEST is structurally conserved with epoxide hydrolases, a class of proteins involved in the biosynthesis of specialized proresolving mediators (SPMs)^47, 48^. SPMs are signaling molecules in the resolution of inflammation and include maresins, resolvins, lipoxins, and protectins, which broadly aid in tissue regeneration^46, 49^. Taken together, these data suggest Mest is critical for resolution of inflammation and moving forward it will be important to experimentally anchor MEST to this process.

A key finding in this study is that regulation of Mest transcription in the digit tip blastema can escape genomic imprinting via promoter switching. A priori, we anticipated persistence of maternal imprinting in the blastema as has been defined for Mest during embryogenesis and in many adult tissues^7, 16^. Consistent with this hypothesis, we found that the canonical variant Mest211, regulated by the DMR surrounding the conical promoter, contributed to the majority of Mest expression in Mest-WT and mKO blastemas (Figures 4E, G, 5B, C). Moreover, no significant expression of any Mest transcript variant was found in Mest-KO blastemas (Figure 4C), as would be expected for a genetic knockout. However, while Mest-pKO blastemas maintain maternal imprinting of the Mest canonical promoter and have negligible Mest211 expression, Mest expression was unexpectedly found in the form of the Mest210 variant (Figure 4E, G, 5D, E). This aligns with altered regulation via the upstream promoter, whereby Mest-pKO and KO blastemas have decreased methylation of the CTCF consensus region (Figure 4I), which is associated with increased CTCF binding to modulate DNA conformational changes^50, 51^. While we found increased methylation surrounding exon A (Figure 4K), this region can be defined as a low CpG promoter (five CpG in 245bp, Figure S6C,E), in which DNA methylation has been shown to increase transcription in various contexts^52–54^.

These data provide strong evidence that reduced expression of Mest during digit tip regeneration results in Mest upregulation via promoter switching, not loss of imprinting. Why there is a demand to maintain high levels of Mest in the blastema remains to be understood, though it is clear this is not simply an artifact of our genetic system. We find almost no Mest210 transcription in e12.5 embryos and no significant change in the canonical or alternative Mest promoter between Mest-WT and pKO embryos (Figures 4, 5), indicating there is no compensation for genetic loss of Mest during embryogenesis. However, promoter switching^30, 35^ and biallelic expression at the Mest locus have been reported, primarily in human cancers^20–22, 30^. The mechanism behind the promoter switching, and whether a discrete set of transcription factors is required, is not yet known. While our study focuses on the regulation and function of a single gene, the connection we find between the blastema and cancer is a classical concept that has not been experimentally resolved^55, 56^. Our study shows that Mest is poised to shed light on this topic, and in future work it will be important to determine how many genes in the blastema share regulatory mechanisms with cancers when under genetic pressure.

## MATERIALS AND METHODS

### Mice

All mice were housed in the Brigham and Women’s Hospital Hale BTM vivarium, maintained by the Center for Comparative Medicine. Breeding and surgery protocols were approved by BWH IACUC. Male and female wildtype 8-week-old FVB/NJ mice (JAX #001800) were used for HCR FISH expression analyses and 28dpa scRNAseq. Mest genetic analyses used the existing Mest knockout allele, B6-Mest^tm1.2Rkz^ (courtesy of Dr. Robert Koza, MMCRI)^29^. Mest allele genotyping primers from the original publication were used (Table S1)^29^.

### Mouse digit tip amputation surgeries

Male and female 8 to 10-week-old mice were used for digit amputation surgeries and subsequent tissue collection. Mice were anesthetized with inhaled 1-2% isoflurane in oxygen during surgeries. For both hindlimbs, digits 2, 3, and 4 were amputated using a sterile disposable scalpel. Distal (P3), regenerative amputations were made midway through the third phalangeal bone; proximal (P2), nonregenerative amputations were made midway through the second phalangeal bone. For analgesia, subcutaneous buprenorphine HCl (0.05mg/kg) was given peri- and post-surgically. Mice were euthanized by carbon dioxide inhalation for post-amputation tissue collections.

### Hybridization Chain Reaction RNA Fluorescent in Situ Hybridization (HCR-FISH)

Post-amputation digits were fixed in 4% PFA at 4°C overnight, followed by PBS washing and decalcification (Decalcifying Solution-Lite, Sigma-Aldrich). Tissues were cryopreserved through a sucrose gradient, embedded in OCT (Tissue-Tek), and sectioned at 20µm with a Leica CM3050s cryostat. HCR-FISH protocol was performed as previously reported^3, 26^, using a commercial Mest probe (Genbank: NM_001252292.1; Molecular Instruments), and were counterstained with 1ng/µL DAPI. TrueVIEW autofluoresence quenching kit (Vector Laboratories) was used for background reduction. Slides were imaged on a Zeiss LSM880 confocal microscope and z-stack images were processed using Fiji^57^.

### Masson Trichrome histology and Alizarin Red bone staining

For Masson Trichrome histology, post-amputation digits were fixed in 4% PFA at 4°C overnight, decalcified with Surgipath Decalcifier I (Leica Biosystems), and cryopreserved through a sucrose gradient to OCT (Tissue-Tek). 14µm sections made with a Leica CM3050s cryostat were stained using a Masson Trichrome Staining Kit (Sigma-Aldrich). Manufacturer’s protocol was followed except for the omission of deparaffinization and microwaving. Stained sections were imaged with a Leica DM2000 upright microscope and images were processed and analyzed with Fiji^57^. 2D blastema area was quantified using Fiji and one-way ANOVA followed by post-hoc t-test with Bonferroni correction was used to assess statistical significance of the measurements among genotypes.

Whole mount Alizarin Red stained digit tip bones were generated as previously reported^58^. Post-amputation digits and contralateral unamputated control digits were collected for skeletal staining at 28dpa and 56dpa; 6-8 mice were used for each genotype. Stained bones were imaged on a Leica M165FC stereomicroscope using the LAS X imaging software. Images were analyzed in Fiji to measure the 2D area and length of the P3 bone. Length was measured from the midpoint at the base of the P3 bone to the tip of the bone. Percent regeneration was calculated for each regenerated digit as 2D area normalized to the average area of the unamputated digits for its respective genotype. Statistical analysis among the genotypes was performed with one-way ANOVA followed by post-hoc t-test with Bonferroni correction.

### Cell culture and in vitro MSC differentiation

To generate the Mest overexpression plasmid, we utilized the pCAG-GFP vector^59^ (pCAG-GFP was a gift from Connie Cepko; Addgene #11150). GFP was excised with EcoRI and NotI digestion and a synthesized Mest211 cDNA gBlock (Genbank: NM_008590; Integrated DNA Technologies) was inserted with NEBuilder HiFi DNA assembly (New England Biolabs). The final construct was verified by plasmid DNA sequencing (Plasmidsaurus).

Mouse multipotent C3H10T1/2 cells (ATCC #CCL-226) and mouse pre-osteoblastic MC3T3-E1 cells (ATCC #CRL-2593) were maintained in αMEM media with 2mM L-glutamine, 1% penicillin-streptomycin and 10% FBS. 20,000 cells were seeded in 12-well plates and transfected with 300ng of pCAG-Mest or pCAG-GFP control plasmid using FuGENE HD transfection reagent (Promega) per manufacturer’s protocol. Lineage-appropriate differentiation media was added 24 hours post-transfection. For osteodifferentiation, media composition was αMEM with 10% FBS, 1% penicillin-streptomycin, 10mM β-glycerophosphate, 50ug/mL ascorbic acid, 100nM dexamethasone^60^. Cells were fixed with 4% PFA and stained with 40mM Alizarin Red to visualize mineralization which was quantified using Fiji^57^. For adipogenic differentiation, media was composed of DMEM with 10% FBS, 1% penicillin-streptomycin, 100nM dexamethasone, 0.5µM isobutylmethylxanthine, and 10ng/mL insulin^61^. Terminal differentiation was assessed by Oil Red O staining which was quantified by OD490nm reading of isopropanol extracted dye. For chondrogenic differentiation, tissue culture plates coated with 10% gelatin were used. Media was composed of DMEM with 10% FBS, 1% penicillin-streptomycin, 100nM dexamethasone, 1% insulin-transferrin-selenium, 50µM L-ascorbic acid 2-phosphate, 50µg/mL proline, and 20ng/mL TGFβ3^61^. Differentiated chondrocytes were visualized with Alcian Blue staining which was quantified by OD595nm reading of hydrochloric acid extracted dye.

### Protein purification and western blot

For validation of pCAG-Mest construct expression, cells were harvested 2 days post transfection in RIPA buffer with protease inhibitor. 30µg of total protein lysate was loaded onto 4-20% Tris-Glycine polyacrylamide gels and transferred onto 45µm nitrocellulose membrane. Primary antibodies were anti-β-tubulin (1:1000, ThermoFisher 32-2600) and anti-Mest (1:1000, Abcam ab151564) used with HRP conjugated secondary antibodies (Jackson ImmunoResearch).

### RNA purification and qPCR

For in vitro experiments, cells were collected from plates on the day noted and RNA was purified using the RNeasy mini prep kit (Qiagen), following the manufacturer’s protocol. For mouse experiments, 30-48 12dpa blastemas from each genotype were dissected and pooled. Timed pregnant mice were used to collect e12.5 Mest-WT and Mest-pKO embryos. RNA extraction for blastema and embryo tissues was performed using TRIzol reagent (Invitrogen), following the manufacturer’s protocol; tissues were homogenized using Navy Eppendorf RNA lysis tubes with the Bullet Blender Storm24 (Next Advance) at 4°C. cDNA was synthesized with SuperScript IV First Strand Synthesis kit (Invitrogen) primed with oligo-dT. Quantitative PCRs for Perilipin-1, PPARγ, Col2a1, Col10a1, Runx2, Mest(total), Mest202, Mest210, and Mest211 were performed using SsoAdvanced Universal SYBR green supermix (BioRad) on QuantStudio 5 Real-time PCR machine. ΔCt was calculated for each well using GAPDH as a housekeeping gene. In-plate technical triplicates allowed for calculation of average relative expression using the 2^(-ΔCt)^ method^62^. All qPCR experiments were performed in triplicate; qPCR primers used for all genes, including those previously published^29, 61, 63–65^, are detailed in Table S1. For miRNA expression, gene-specific cDNA was synthesized using U6 and miR-335 primers with the Taqman MicroRNA Reverse Transcription kit (ThermoFisher). qPCR was performed using U6 and miR-335 TaqMan assays (001973, 000546; ThermoFisher). Statistical analyses were performed by two-sample t-test, one-way ANOVA followed by post-hoc t-tests with Bonferroni correction, or two-way ANOVA followed by post-hoc t-tests with Tukey HSD correction.

### Bisulfite sequencing

12-18 12dpa blastemas were collected and pooled for each Mest genotype group and hindlimbs were dissected from two e12.5 Mest-WT and Mest-pKO embryos. Genomic DNA was purified with the DNeasy Blood and Tissue Kit (Qiagen). 500ng of genomic DNA was used for bisulfite conversion with the EZ DNA Methylation Kit (Zymo Research), following the manufacturer’s protocol. Bisulfite converted DNA was used for PCR amplification for the four regions of interest: CTCF, ExonA, Exon1-5’, and Exon1-3’, using ZymoTaq DNA polymerase (Zymo Research). Primers used for amplification were designed with the Bisulfite Primer Design Tool (www.zymoresearch.com/pages/bisulfite-primer-seeker) and MethPrimer (www.urogene.org/methprimer/) (Table S1). PCR amplicons were TA cloned into pGEM-T-Easy (Promega) to facilitate allele-specific DNA sequencing from the Sp6 promoter (Eton Biosciences). A minimum of 10 clones were sequenced per genotype for each genomic region. Each sequencing read was scored for number of cytosines present in the appropriate amplicon region and compared to the predicted, fully methylated bisulfite converted sequence. Thus, an observed:predicted cytosine ratio of 0 indicates hypomethylation while a ratio of 1 indicates hypermethylation. Statistical analysis was performed using the Chi-square discrete test for uniformity, using the number of predicted methylated cytosines as the denominator for the test. One-way ANOVA followed by post-hoc t-tests with Bonferroni correction was used for statistical analysis.

### Single cell isolation, library construction and RNA sequencing

All single cell transcriptomics were performed using the 10X Chromium single cell gene expression platform (10X Genomics) at the BWH Center for Cellular Profiling. The 28dpa single cell isolation, library construction, and sequencing were performed as previously published reported^3^. For Mest-WT and Mest-KO scRNAseq, 36 12dpa blastemas were dissected and pooled for each genotype. To minimize any potential batch effect, these were isolated and underwent library construction on the same day. Single-cell suspensions were generated for each genotype as previously described^3^, with the addition of FACS for live cells using propidium iodide. In short, tissue was digested with 2.5% trypsin-EDTA and 10% collagenase, followed by mechanical dissociation by pipetting. Cells were strained through 35µm filters and red blood cells were removed with ACK lysing buffer (Gibco). Cells were resuspended in 0.4% BSA/PBS and 1µg/µL propidium iodide. 10,000 propidium iodide negative cells were collected into 0.4% BSA/PBS and immediately used for 10x Genomics droplet encapsulation. cDNA libraries were made using the Single Cell 3’ v3 chemistry (10x Genomics). All libraries were sequenced at the DFCI Molecular Biology Core Facility by Illumina NextSeq 500.

For neutrophil flow cytometry, Mest-WT and KO regenerating digit tips were harvested at 9, 12, and 15dpa. Single-cell suspensions were generated as described above with an additional incubation with anti-CD11b-BV421 (1:200, Biolegend #101251) and anti-Ly6g-AF647 (1:200, Biolegend #127609) prior to the addition of propidium iodide. Samples were sorted on a BD FACSAria, gating to remove doublets based on FSC-A and FSC-H, isolating PI^-^/CD11b^+^/Ly6g^+^ cells. Analysis was performed using FloJo version 10.8.1 (BD Biosciences). Percentages of double positive CD11b and Ly6g of all PI^-^ cells were calculated for each genotype and time point. Cell numbers for PI^-^ and CD11b^+^/Ly6g^+^ cells were used for differential proportion analysis between Mest-WT and KO at each time point.

### scRNAseq computational analyses

High-performance computing was done with the O2 cluster supported by the Harvard Medical School Research Computing Group (it.hms.harvard.edu/service/high-performance-computing-hpc). FASTQ files for all scRNAseq experiments were aligned to the mouse mm10-2020-A transcriptome with introns using CellRanger version 7.1.0 (10x Genomics). Cell by gene matrices for each dataset were individually imported into R version 4.3.1^66^ for analysis using Seurat version 5.0.1^67^. Cells were filtered based on percent mitochondrial genes, UMI count, and doublet prediction as previously reported^3^. Post-filtering, the UA, 11, 12, 14, 17, and 28dpa datasets contained 11499, 6918, 3039, 5414, 7947, and 8591 singlets, respectively, which we utilized for further analyses. The Mest-WT and Mest-KO 12dpa datasets contained 8805 and 6898 singlets, respectively, used for analyses. All individual datasets were processed via normalization, identification of variable features, scaling, linear dimensional reduction, and clustering as previously described^3^.

Integration of the UA, 11, 12, 14, 17, and 28dpa data was performed using the CCA Integration method in Seurat followed by re-normalization, scaling, and clustering. Fibroblast clusters were identified and subsetted by high expression of marker genes Prx1, Pdgfrα, and Lum. These subsetted fibroblasts were used for lineage trajectory, using the web interface (https://kleintools.hms.harvard.edu/tools/spring.html) for SPRING^15^ analysis. Integration of Mest-WT and Mest-KO datasets was performed using the CCA Integration method in Seurat followed by re-normalization, scaling, and clustering. Fibroblast clusters were identified and subsetted based on high expression of Pdgfrα and Lum. Immune cell clusters were identified and subsetted based on expression of Lyz2 (macrophages/monocytes), Cd3g (T-cells), Ctsk (pre-osteoclasts), and S100a9 (neutrophils). Differentially expressed genes (DEGs) for fibroblast subpopulations and between Mest genotypes were determined using Seurat FindAllMarkers and FindMarkers, respectively, with the following parameters: min.pct=0.25 and avg.log2FC >1.00 (for fibroblast cluster DEGs) or avg.log2FC > 0.58 (for Mest-WT and KO DEGs) and adj.pval < 0.05 (Tables S2-S4). ClusterProfiler version 4.10.0 was used to perform gene set enrichment analysis of DEGs with KEGG pathway^68^. To determine statistically significant changes in populations between Mest-WT and Mest-KO cell types, fibroblast and immune cell clusters, we utilized differential proportion analysis as previously reported^3, 69^. Signaling changes were assessed using our defined cell types with CellChat version 2.1.1^40^ and NICHES version 1.0.0^41^, utilizing the mouse CellChatDB and omnipath ligand-receptor databases, respectively.

## Supporting information

supplement_all

## DATA AVAILABILITY

All raw single cell RNAseq FASTQ files and cell by gene expression matrices are available via NCBI GEO: GSE143888, GSE267446, and GSE267553.

## ACKNOWLEDGEMENTS

We thank Affan Shaikh and Scott Semelsberger for their assistance with technical troubleshooting, and Amanda Whipple and Bongmin Bae for helpful conversations. We thank Robert Koza (Maine Medical Center) for sharing the Mest knockout mouse allele. We acknowledge the NeuroTechnology Studio at Brigham and Women’s Hospital for providing Zeiss LSM880 confocal microscope access and consultation on data acquisition. This work was supported by the Eunice Kennedy Shriver National Institute of Child and Human Development (R01HD109200 to J.A.L) and the Harvard Stem Cell Institute (DP-0205-2201 to J.A.L). V.J received support from NIH/NIAMS T32AR055885 and F31AR082220.

## AUTHOR CONTRIBUTIONS

V.J and J.A.L conceptualized and designed the study. V.J and S.M.P performed the experiments. V.J and J.A.L performed the computational analyses. V.J, S.M.P, and J.A.L analyzed and interpreted the data. V.J and J.A.L wrote the manuscript, and all authors approved the final version.

