## supplement_all for "Regeneration-specific promoter switching facilitates Mest expression in the mouse digit tip to modulate neutrophil response"

### SUPPLEMENTAL FIGURES

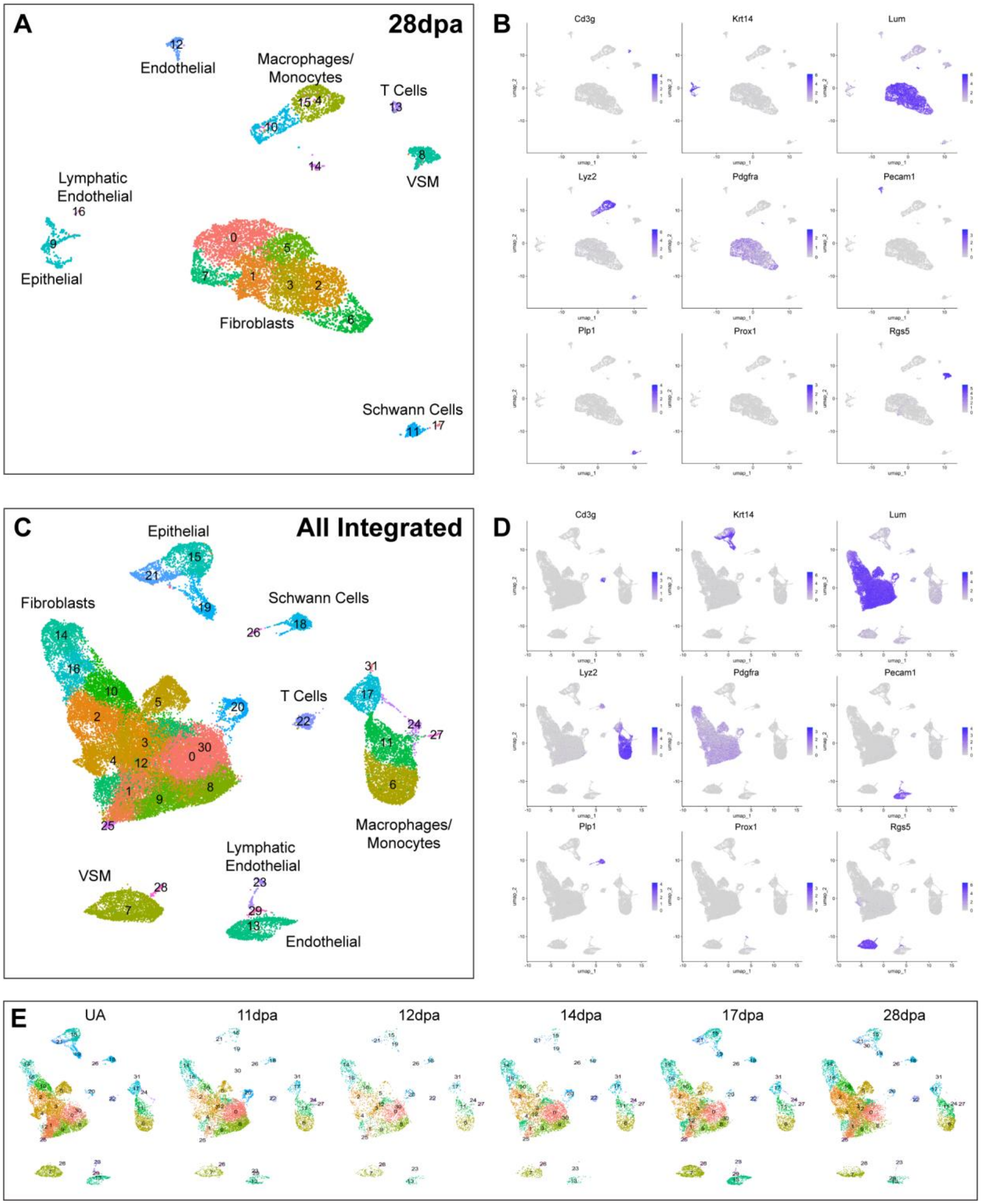

**Supplemental Figure 1**

**Longitudinal regenerating digit tip scRNAseq analysis including 28dpa.** (A) UMAP plot of cell clustering from 28dpa regenerated digit tip scRNAseq; (B) gene markers used to assign cluster identities. (C) UMAP plot of cell clustering from integrated UA, 11, 12, 14, 17, and 28dpa scRNAseq datasets; (D) gene markers used to assign cluster identities. (E) UMAP plots of integrated scRNAseq datasets, split by stage.

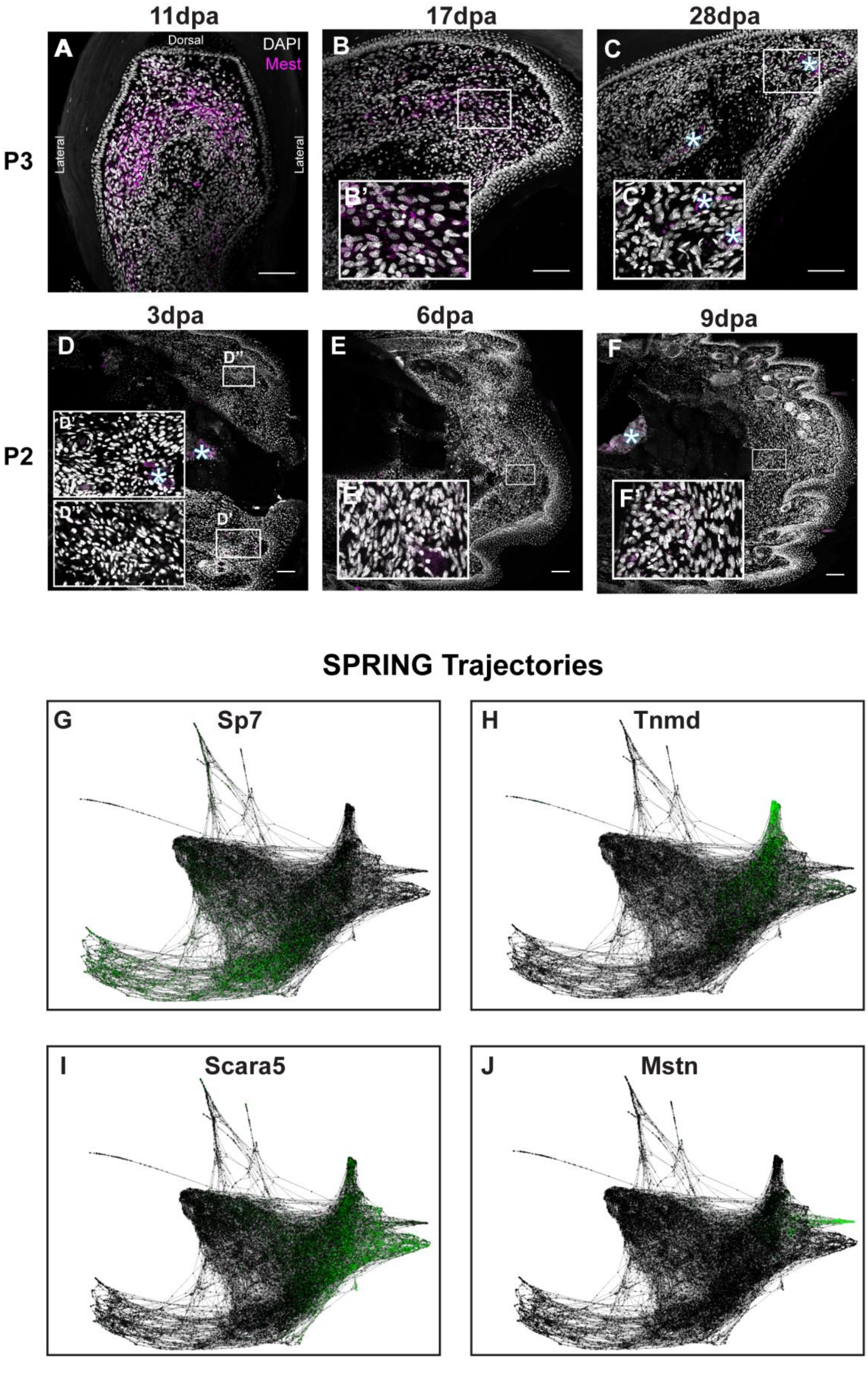
11dpa
17dpa
28dpa
A
Dorsal
DAPI
Mest
Lateral
Lateral
B
B'
C
C'
P3
3dpa
6dpa
9dpa
D
D''
D'
D'
D''
E
E'
F
F'
P2
SPRING Trajectories
G
Sp7
H
Tnmd
I
Scara5
J
Mstn

**Supplemental Figure 2**

**Post-amputation Mest expression in distal P3 and proximal P2 digits.** Mest RNA expression (magenta) in post-amputation digit sections visualized by HCR-RNA FISH, counterstained with DAPI (grayscale). (A-C) Regenerating P3 digits at (A) 11dpa, (B) 17dpa, and (C) 28dpa. (D-F) Mest expression in the P2 non-regenerating digit at (D) 3dpa, (E) 6dpa, and (F) 9dpa. The tissue orientation is distal to the right and dorsal to the top for all panels except (A) where dorsal is to the top and lateral is to the right and left. White boxes show locations of inset panels (B' - F'); all scale bars = 100µm. White asterisks denote blood vessel autofluorescence. (G-J) SPRING plots of integrated fibroblasts with expression of the genes (G) Sp7 (osteogenic), (H) Tnmd (ligaments and tendons), (I) Scara5 (dermal), and (J) Mstn (myogenic and connective tissue) in green.

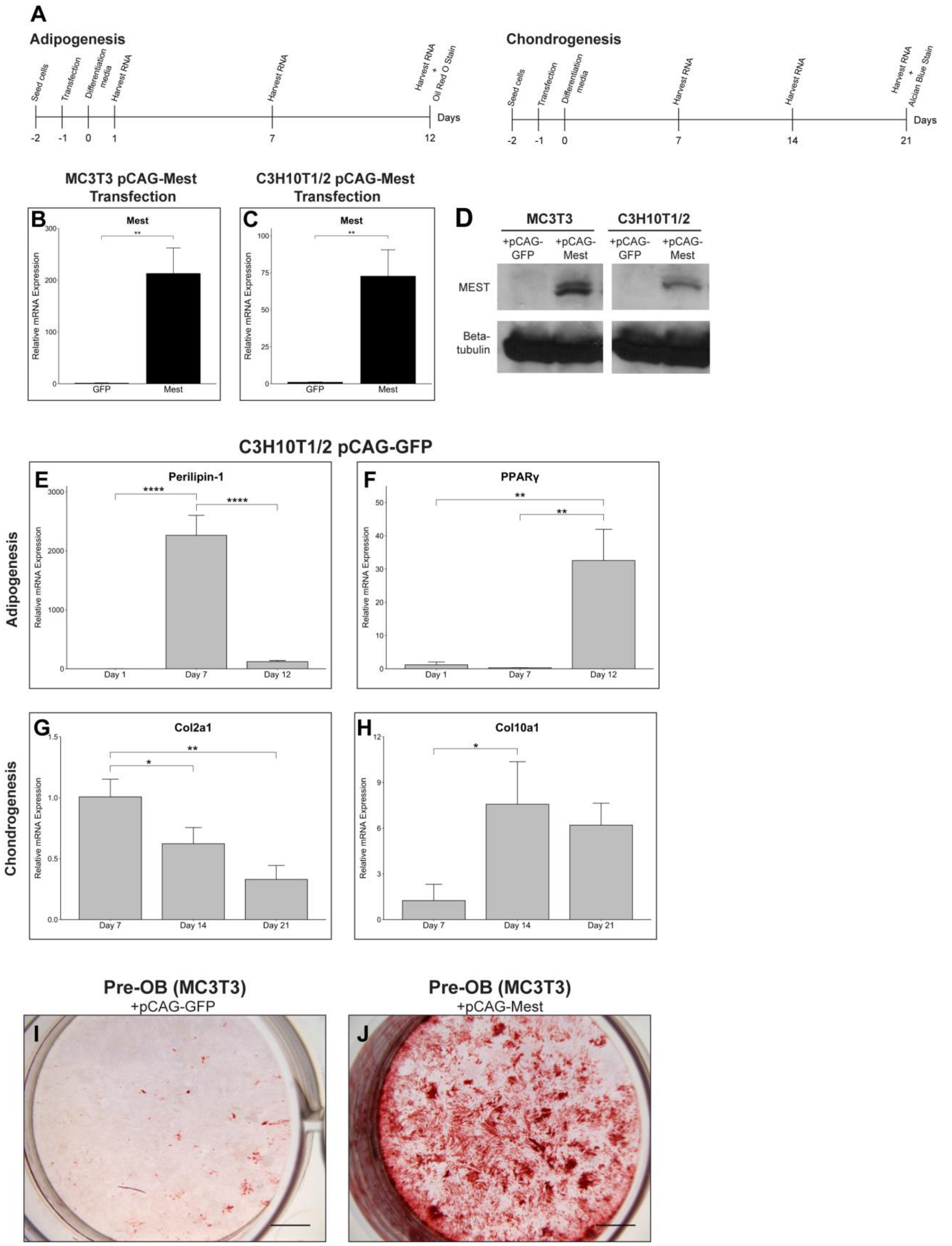

A
Adipogenesis
Seed cells
Transfection
Differentiation media
Harvest RNA
Harvest RNA
Harvest RNA + Oil Red O Stain
Days
-2 -1 0 1 7 12
Chondrogenesis
Seed cells
Transfection
Differentiation media
Harvest RNA
Harvest RNA
Harvest RNA + Alcian Blue Stain
Days
-2 -1 0 7 14 21
MC3T3 pCAG-Mest Transfection
B
Mest
Relative mRNA Expression
GFP
Mest
C3H10T1/2 pCAG-Mest Transfection
C
Mest
Relative mRNA Expression
GFP
Mest
D
MC3T3
C3H10T1/2
+pCAG-GFP
+pCAG-Mest
+pCAG-GFP
+pCAG-Mest
MEST
Beta-tubulin
C3H10T1/2 pCAG-GFP
Adipogenesis
E
Perilipin-1
Relative mRNA Expression
Day 1
Day 7
Day 12
F
PPARγ
Relative mRNA Expression
Day 1
Day 7
Day 12
Chondrogenesis
G
Col2a1
Relative mRNA Expression
Day 7
Day 14
Day 21
H
Col10a1
Relative mRNA Expression
Day 7
Day 14
Day 21
Pre-OB (MC3T3)
+pCAG-GFP
I
Pre-OB (MC3T3)
+pCAG-Mest
J

**Supplemental Figure 3**

**In vitro Mest overexpression experiments.** (A) In vitro (left) adipogenesis and (right) chondrogenesis experimental timelines. (B-C) Mest qPCR analysis of pCAG-GFP and pCAG-Mest transfected (B) MC3T3 and (C) C3H10T1/2 cells 24 hours post-transfection. (D) MEST protein and Beta-tubulin control western blot analysis of pCAG-GFP and pCAG-Mest transfected (left) MC3T3 and (right) C3H10T1/2 cells 48 hours post transfection. (E-F) qPCR analysis of adipogenic genes (E) Perilipin-1 and (F) PPARγ of 1, 7, and 12dpi C3H10T1/2 cells transfected with GFP. (G-H) qPCR analysis of chondrogenic genes (G) Col2a1 and (H) Col10a1 of 7, 14, and 21dpi C3H10T1/2 cells transfected with GFP. (I-J) Alizarin Red stained wells of 28dpi (I) GFP and (J) Mest transfected MC3T3 cells for osteogenesis. Scale bars = 250µm. Error bars are reported as standard deviation; (*) $p < 0.05$, (**) $p < 0.01$, (****) $p < 0.0001$.

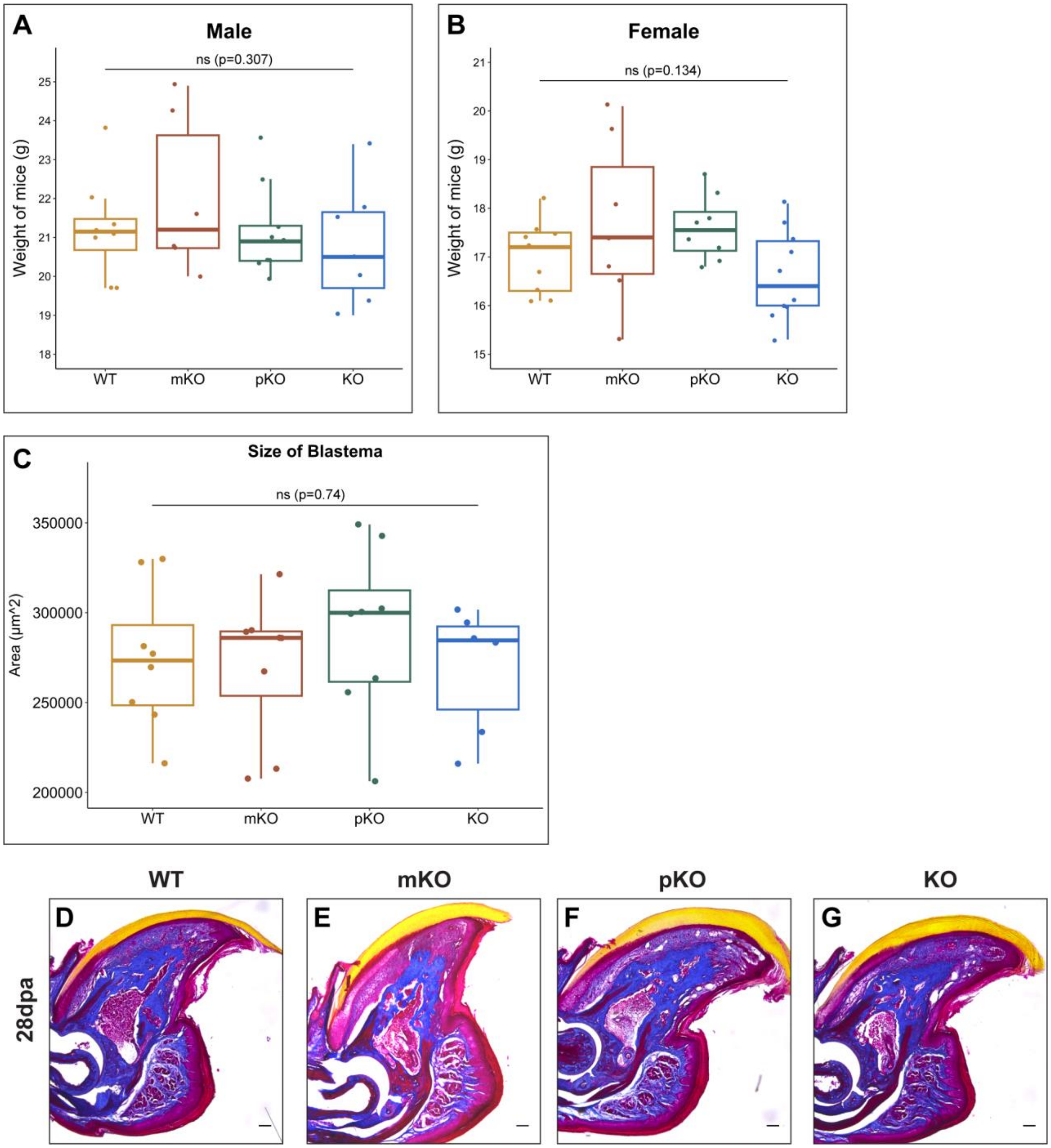

A
Male
ns (p=0.307)
Weight of mice (g)
WT
mKO
pKO
KO
B
Female
ns (p=0.134)
Weight of mice (g)
WT
mKO
pKO
KO
C
Size of Blastema
ns (p=0.74)
Area (µm^2)
WT
mKO
pKO
KO
WT
mKO
pKO
KO
D
E
F
G
28dpa

**Supplemental Figure 4**

**Extended data for Mest genetic cohorts.** (A, B) Boxplot of weights of 8-10-week-old Mest knockout mice for (A) males and (B) females. (C) Boxplot of 12dpa blastema sizes across Mest genetic cohorts. All data was analyzed with one-way ANOVA and was not statistically significant (ns) for either the male or female cohorts for weight and for the size of blastemas ($p=0.307$, 0.134, and 0.740, respectively). (D-G) Masson trichrome histology of 28dpa regenerated digit sections from (D) WT, (E) mKO, (F) pKO, and (G) KO mice. Scale bars = 100µm.

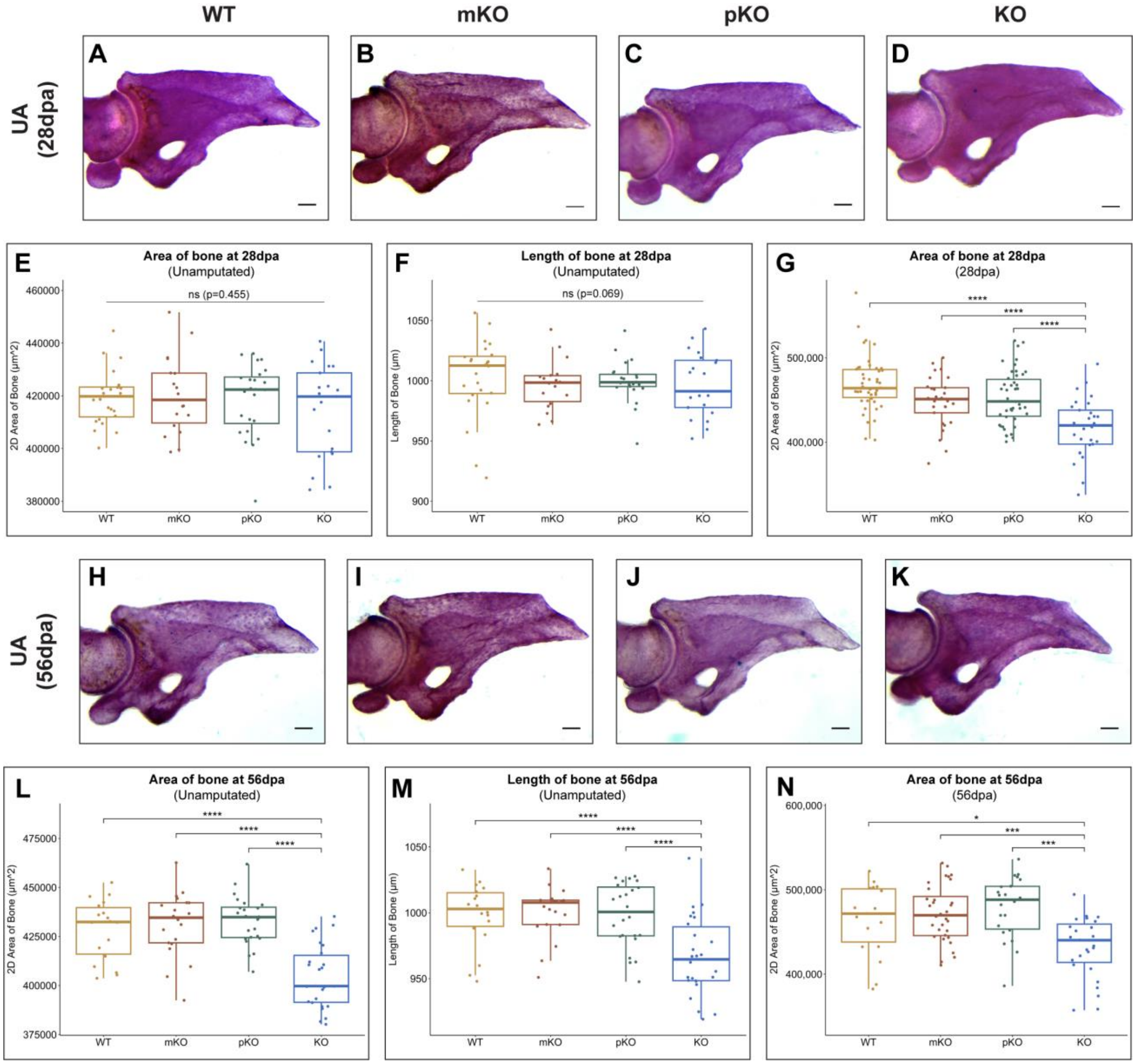

WT
mKO
pKO
KO
UA (28dpa)
A
B
C
D
E
Area of bone at 28dpa
(Unamputated)
ns (p=0.455)
2D Area of Bone (μm^2)
460000
440000
420000
400000
380000
WT
mKO
pKO
KO
F
Length of bone at 28dpa
(Unamputated)
ns (p=0.069)
Length of Bone (μm)
1050
1000
950
900
WT
mKO
pKO
KO
G
Area of bone at 28dpa
(28dpa)
****
****
****
2D Area of Bone (μm^2)
500,000
400,000
WT
mKO
pKO
KO
UA (56dpa)
H
I
J
K
L
Area of bone at 56dpa
(Unamputated)
****
****
****
2D Area of Bone (μm^2)
475000
450000
425000
400000
375000
WT
mKO
pKO
KO
M
Length of bone at 56dpa
(Unamputated)
****
****
****
Length of Bone (μm)
1050
1000
950
WT
mKO
pKO
KO
N
Area of bone at 56dpa
(56dpa)
*
***
***
2D Area of Bone (μm^2)
600,000
500,000
400,000
WT
mKO
pKO
KO

**Supplemental Figure 5**

**Extended data for Alizarin Red stained whole-mount digits.** (A-D) Representative Alizarin Red stained unamputated contralateral digits for Mest genetic cohorts harvested at 28dpa. (E, F) Quantification of (E) 2D area of bone and (F) length of the P3 bone for unamputated, 28dpa digits. Data were not significant (ns) by one-way ANOVA ($p=0.455$ and 0.0694, respectively). (G) Quantification of 2D area for amputated bones at 28dpa. (H-K) Representative Alizarin Red stained unamputated contralateral digits for Mest cohorts harvested at 56dpa. (L, M) Quantification of (L) 2D area of bone and (M) length of the P3 bone for the unamputated, 56dpa digits. (N) 2D area of bone of amputated bones at 56dpa. All data was analyzed with one-way ANOVA followed by post-hoc t-test with Bonferroni correction. (*) $p < 0.05$, (***) $p < 0.001$, (****) $p < 0.0001$. Scale bars = 100μm.

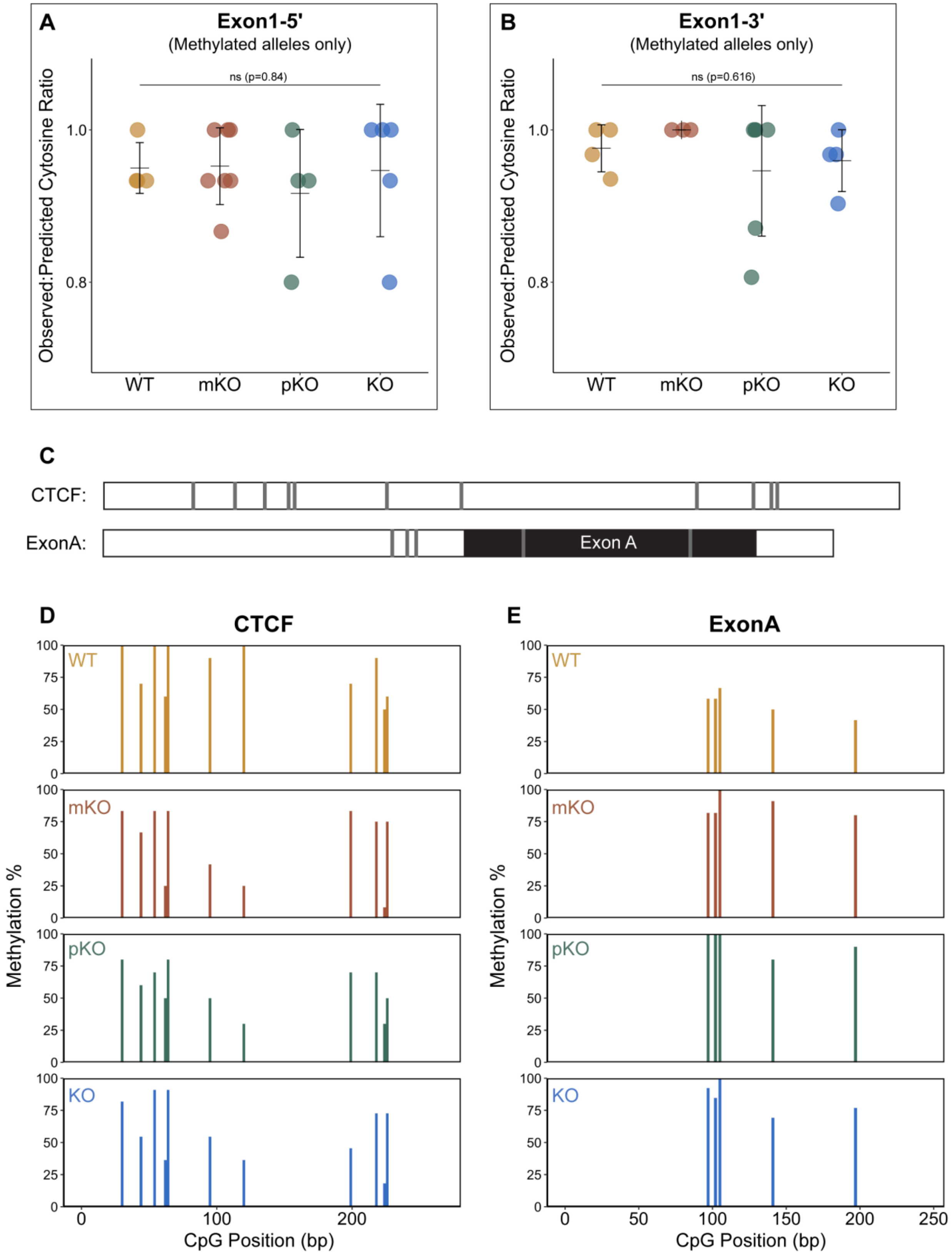

A
Exon1-5'
(Methylated alleles only)
ns (p=0.84)
Observed:Predicted Cytosine Ratio
1.0
0.8
WT
mKO
pKO
KO
B
Exon1-3'
(Methylated alleles only)
ns (p=0.616)
Observed:Predicted Cytosine Ratio
1.0
0.8
WT
mKO
pKO
KO
C
CTCF:
ExonA:
Exon A
D
CTCF
E
ExonA
WT
mKO
pKO
KO
Methylation %
100
75
50
25
0
0
50
100
150
200
250
CpG Position (bp)

**Supplemental Figure 6**

**Extended data for bisulfite sequencing of Mest cohorts.** (A-B) Dot plots of methylated alleles only in (A) Exon1-5' and (B) Exon1-3'. One-way ANOVA analyzes indicated no significance in both datasets (p=0.84 and 0.616, respectively). (C) Schematic of (top) CTCF and (bottom) ExonA amplicons for bisulfite sequencing. Gray bars indicate predicted methylated CpG position and black box indicates the location of exon A. (D-E) Graphs of methylation percentage at each predicted CpG position across all reads for each genotype in the (D) CTCF and (E) ExonA regions.

**A**

```
Mest211  1    MVRRD-------RLRRMREWWVQVGLLAVPLLAAYLHIPPPQLSPALHSWKTSGKFFTYKGLRIFYQDSVGVVGSPEIVV   73
Mest210  1    ----------------MREWWVQVGLLAVPLLAAYLHIPPPQLSPALHSWKTSGKFFTYKGLRIFYQDSVGVVGSPEIVV   64
Mest202  1    MYVLEprssslLKAMQMREWWVQVGLLAVPLLAAYLHIPPPQLSPALHSWKTSGKFFTYKGLRIFYQDSVGVVGSPEIVV   80

Mest211  74   LLHGFPTSSYDWYKIWEGLTLRFHRVIALDFLGFGFSDKPRPHQYSIFEQASIVESLLRHLGLQNRRINLLSHDYGDIVA   153
Mest210  65   LLHGFPTSSYDWYKIWEGLTLRFHRVIALDFLGFGFSDKPRPHQYSIFEQASIVESLLRHLGLQNRRINLLSHDYGDIVA   144
Mest202  81   LLHGFPTSSYDWYKIWEGLTLRFHRVIALDFLGFGFSDKPRPHQYSIFEQASIVESLLRHLGLQNRRINLLSHDYGDIVA   160

Mest211  154  QELLYRYKQNRSGRLTIKSLCLSNGGIFPETHRPLLLQKLLKDGGVLSPILTRLMNFFVFSRGLTPVFGPYTRPTESELW   233
Mest210  145  QELLYRYKQNRSGRLTIKSLCLSNGGIFPETHRPLLLQKLLKDGGVLSPILTRLMNFFVFSRGLTPVFGPYTRPTESELW   224
Mest202  161  QELLYRYKQNRSGRLTIKSLCLSNGGIFPETHRPLLLQKLLKDGGVLSPILTRLMNFFVFSRGLTPVFGPYTRPTESELW   240

Mest211  234  DMWAVIRNNDGNLVIDSLLQYINQRKKFRRRWVGALASVSIPIHFIYGPLDPINPYPEFLELYRKTLPRSTVSILDDHIS   313
Mest210  225  DMWAVIRNNDGNLVIDSLLQYINQRKKFRRRWVGALASVSIPIHFIYGPLDPINPYPEFLELYRKTLPRSTVSILDDHIS   304
Mest202  241  DMWAVIRNNDGNLVIDSLLQYINQRKKFRRRWVGALASVSIPIHFIYGPLDPINPYPEFLELYRKTLPRSTVSILDDHIS   320

Mest211  314  HYPQLEDPMGFLNAYMGFINSF   335
Mest210  305  HYPQLEDPMGFLNAYMGFINSF   326
Mest202  321  HYPQLEDPMGFLNAYMGFINSF   342
```

**B**

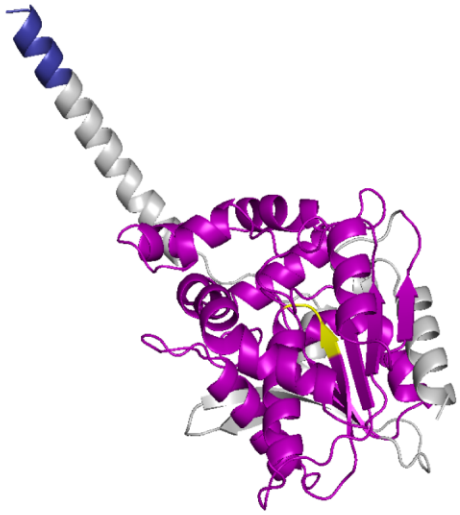

**Dark Blue** = N-terminus region specific to Mest211
**Magenta** = alpha/beta hydrolase domain
**Yellow** = Ser-His-Asp catalytic triad

**Supplemental Figure 7**

**Protein alignment for Mest variants.** (A) Protein sequence alignment for Mest211, 210 and 202 variants with 100% identical amino acids in red font. (B) AlphaFold predicted structure of Mest211 based on #Q07646 with the follow features colored: dark blue = N-terminal amino acids 1-9 specific to Mest211, magenta=alpha/beta hydrolase domain, and yellow = Ser-His-Asp catalytic triad.

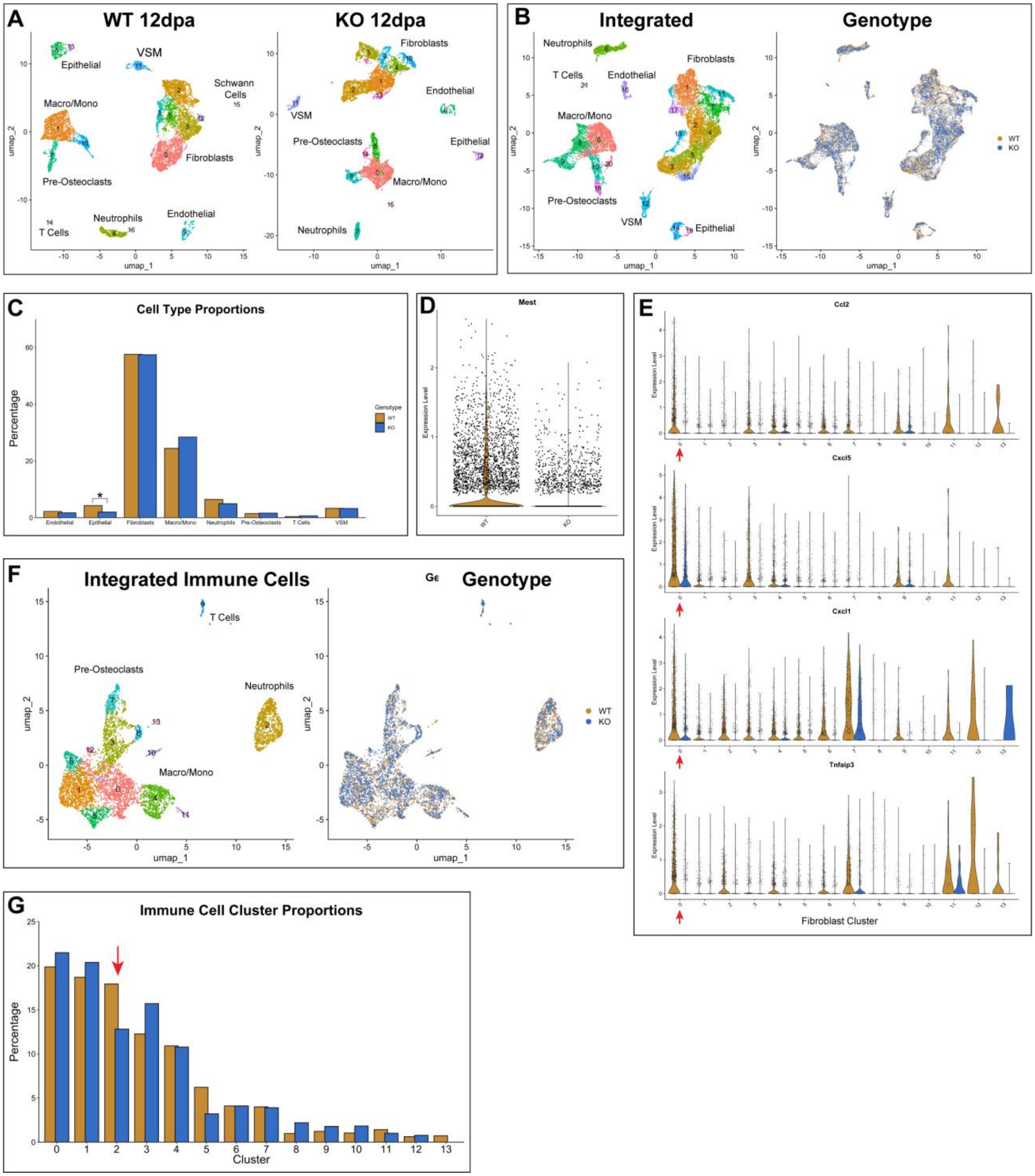
A
WT 12dpa
KO 12dpa
Epithelial
VSM
Schwann Cells
Macro/Mono
Fibroblasts
Pre-Osteoclasts
Neutrophils
Endothelial
T Cells
umap_1
umap_2
B
Integrated
Genotype
WT
KO
C
Cell Type Proportions
Percentage
Genotype
Endothelial
Epithelial
Fibroblasts
Macro/Mono
Neutrophils
Pre-Osteoclasts
T Cells
VSM
D
Mest
Expression Level
E
Ccl2
Cxcl5
Cxcl1
Tnfaip3
Fibroblast Cluster
F
Integrated Immune Cells
Genotype
G
Immune Cell Cluster Proportions
Percentage
Cluster

**Supplemental Figure 8**

**Extended data for Mest scRNAseq.** (A) UMAP plots of (left) Mest-WT and (right) KO 12dpa blastemas colored by Seurat clusters with major cell types labeled. (B) UMAP plot of integrated WT and KO 12dpa blastemas colored by (left) Seurat clusters with major cell types labeled and (right) genotype. (C) Cell type proportions in Mest-WT versus KO datasets. (D) Violin plot of Mest expression in integrated fibroblasts. (E) Expression of Ccl2, Cxcl5, Cxcl1, and Tnfaip3 (the top 4 DEGs in WT fibroblasts) across all fibroblast subpopulations, split by genotype. Cluster 0 is indicated by a red arrow in each graph. (F) UMAP plots of integrated immune cells colored by (left) Seurat clusters and (right) genotype. (G) Cluster proportions in the immune cell subpopulations; red arrow points to cluster 2 neutrophils. Differential proportion analysis was performed for both (C) and (G) and all genotypic comparisons were non-significant, except for the epithelial population in (C).

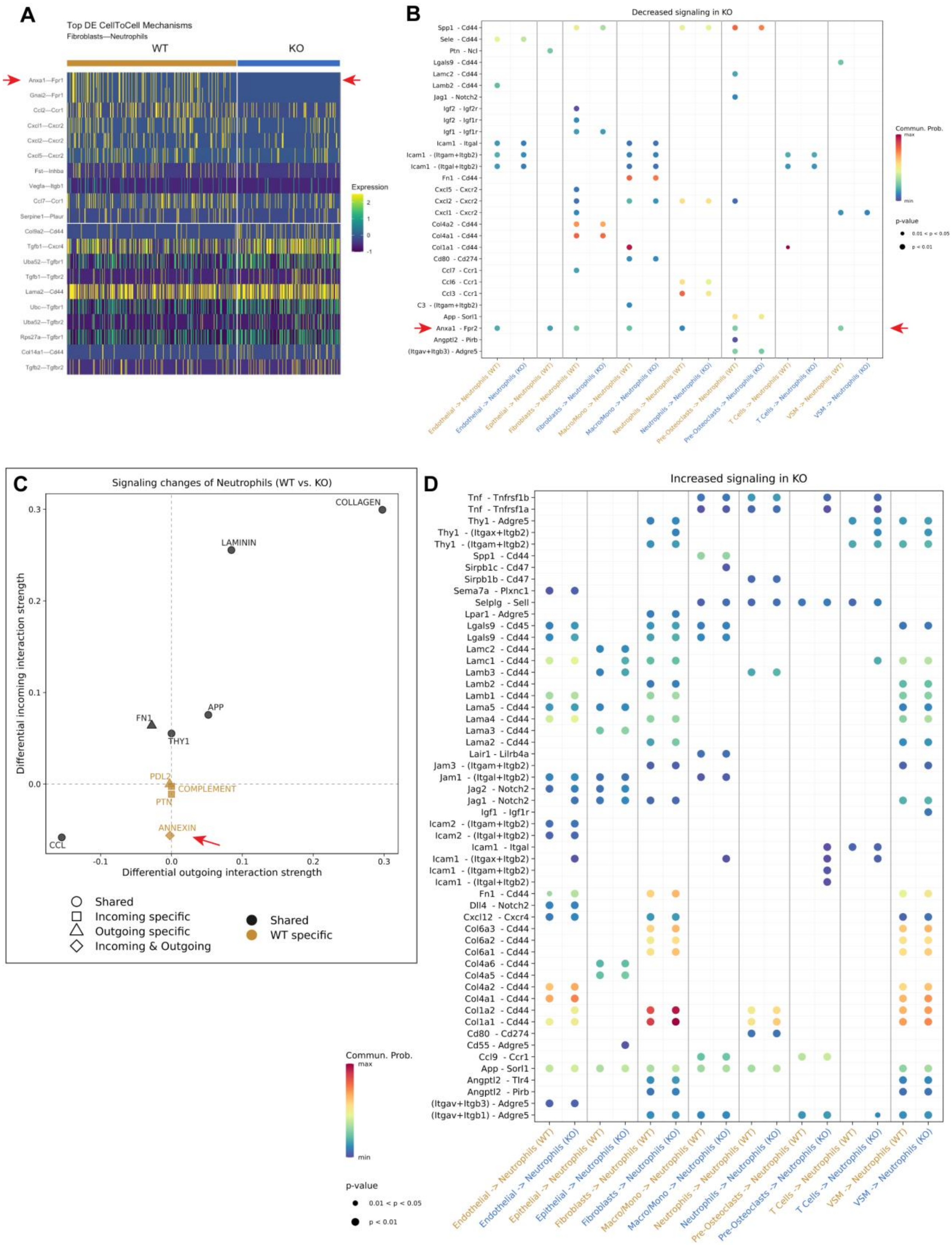
A
Top DE CellToCell Mechanisms
Fibroblasts—Neutrophils
WT
KO
Anxa1—Fpr1
Gnai2—Fpr1
Ccl2—Ccr1
Cxcl1—Cxcr2
Cxcl2—Cxcr2
Cxcl5—Cxcr2
Fst—Inhba
Vegfa—Itgb1
Ccl7—Ccr1
Serpine1—Plaur
Col8a2—Cd44
Tgfb1—Cxcr4
Uba52—Tgfbr1
Tgfb1—Tgfbr2
Lama2—Cd44
Ubc—Tgfbr1
Uba52—Tgfbr2
Rps27a—Tgfbr1
Col14a1—Cd44
Tgfb2—Tgfbr2
Expression
B
Decreased signaling in KO
Spp1 - Cd44
Sele - Cd44
Ptn - Ncl
Lgals9 - Cd44
Lamc2 - Cd44
Lamb2 - Cd44
Jag1 - Notch2
Igf2 - Igf2r
Igf2 - Igf1r
Igf1 - Igf1r
Icam1 - Itgal
Icam1 - (Itgam+Itgb2)
Icam1 - (Itgal+Itgb2)
Fn1 - Cd44
Cxcl5 - Cxcr2
Cxcl2 - Cxcr2
Cxcl1 - Cxcr2
Col4a2 - Cd44
Col4a1 - Cd44
Col1a1 - Cd44
Cd80 - Cd274
Ccl7 - Ccr1
Ccl6 - Ccr1
Ccl3 - Ccr1
C3 - (Itgam+Itgb2)
App - Sorl1
Anxa1 - Fpr2
Angptl2 - Pirb
(Itgav+Itgb3) - Adgre5
Commun. Prob.
max
min
p-value
0.01 < p < 0.05
p < 0.01
Endothelial -> Neutrophils (WT)
Endothelial -> Neutrophils (KO)
Epithelial -> Neutrophils (WT)
Fibroblasts -> Neutrophils (WT)
Fibroblasts -> Neutrophils (KO)
Macro/Mono -> Neutrophils (WT)
Macro/Mono -> Neutrophils (KO)
Neutrophils -> Neutrophils (WT)
Neutrophils -> Neutrophils (KO)
Pre-Osteoclasts -> Neutrophils (WT)
Pre-Osteoclasts -> Neutrophils (KO)
T Cells -> Neutrophils (WT)
T Cells -> Neutrophils (KO)
VSM -> Neutrophils (WT)
VSM -> Neutrophils (KO)
C
Signaling changes of Neutrophils (WT vs. KO)
COLLAGEN
LAMININ
APP
FN1
THY1
PDL2
COMPLEMENT
PTN
ANNEXIN
CCL
Differential outgoing interaction strength
Differential incoming interaction strength
Shared
Incoming specific
Outgoing specific
Incoming & Outgoing
Shared
WT specific
D
Increased signaling in KO
Tnf - Tnfrsf1b
Tnf - Tnfrsf1a
Thy1 - Adgre5
Thy1 - (Itgax+Itgb2)
Thy1 - (Itgam+Itgb2)
Spp1 - Cd44
Sirpb1c - Cd47
Sirpb1b - Cd47
Sema7a - Plxnc1
Selplg - Sell
Lpar1 - Adgre5
Lgals9 - Cd45
Lgals9 - Cd44
Lamc2 - Cd44
Lamc1 - Cd44
Lamb3 - Cd44
Lamb2 - Cd44
Lamb1 - Cd44
Lama5 - Cd44
Lama4 - Cd44
Lama3 - Cd44
Lama2 - Cd44
Lair1 - Lilrb4a
Jam3 - (Itgam+Itgb2)
Jam1 - (Itgal+Itgb2)
Jag2 - Notch2
Jag1 - Notch2
Igf1 - Igf1r
Icam2 - (Itgam+Itgb2)
Icam2 - (Itgal+Itgb2)
Icam1 - Itgal
Icam1 - (Itgax+Itgb2)
Icam1 - (Itgam+Itgb2)
Icam1 - (Itgal+Itgb2)
Fn1 - Cd44
Dll4 - Notch2
Cxcl12 - Cxcr4
Col6a3 - Cd44
Col6a2 - Cd44
Col6a1 - Cd44
Col4a6 - Cd44
Col4a5 - Cd44
Col4a2 - Cd44
Col4a1 - Cd44
Col1a2 - Cd44
Col1a1 - Cd44
Cd80 - Cd274
Cd55 - Adgre5
Ccl9 - Ccr1
App - Sorl1
Angptl2 - Tlr4
Angptl2 - Pirb
(Itgav+Itgb3) - Adgre5
(Itgav+Itgb1) - Adgre5
Commun. Prob.
max
min
p-value
0.01 < p < 0.05
p < 0.01
Endothelial -> Neutrophils (WT)
Endothelial -> Neutrophils (KO)
Epithelial -> Neutrophils (WT)
Epithelial -> Neutrophils (KO)
Fibroblasts -> Neutrophils (WT)
Fibroblasts -> Neutrophils (KO)
Macro/Mono -> Neutrophils (WT)
Macro/Mono -> Neutrophils (KO)
Neutrophils -> Neutrophils (WT)
Neutrophils -> Neutrophils (KO)
Pre-Osteoclasts -> Neutrophils (WT)
Pre-Osteoclasts -> Neutrophils (KO)
T Cells -> Neutrophils (WT)
T Cells -> Neutrophils (KO)
VSM -> Neutrophils (WT)
VSM -> Neutrophils (KO)

**Supplemental Figure 9**

**Extended data for Mest scRNAseq cell-cell communication analysis.** (A) Heatmap of differential ligand-receptor interactions between fibroblasts and neutrophils in the Mest-WT and KO blastemas generated by NICHES analysis. (B, D) Dot plot of ligand-receptor interactions (B) downregulated and (D) upregulated in KO blastemas compared with Mest-WT, with neutrophils as the receiving cell type. (C) Differential strength in signaling pathways with neutrophils as receiving cell type, negative X- and Y-values indicate upregulated in WT blastema while positive X- and Y-values indicate upregulated in KO blastema. Graphs (B-D) were generated using CellChat. Red arrows in (A-C) point to Anxa1 signaling.

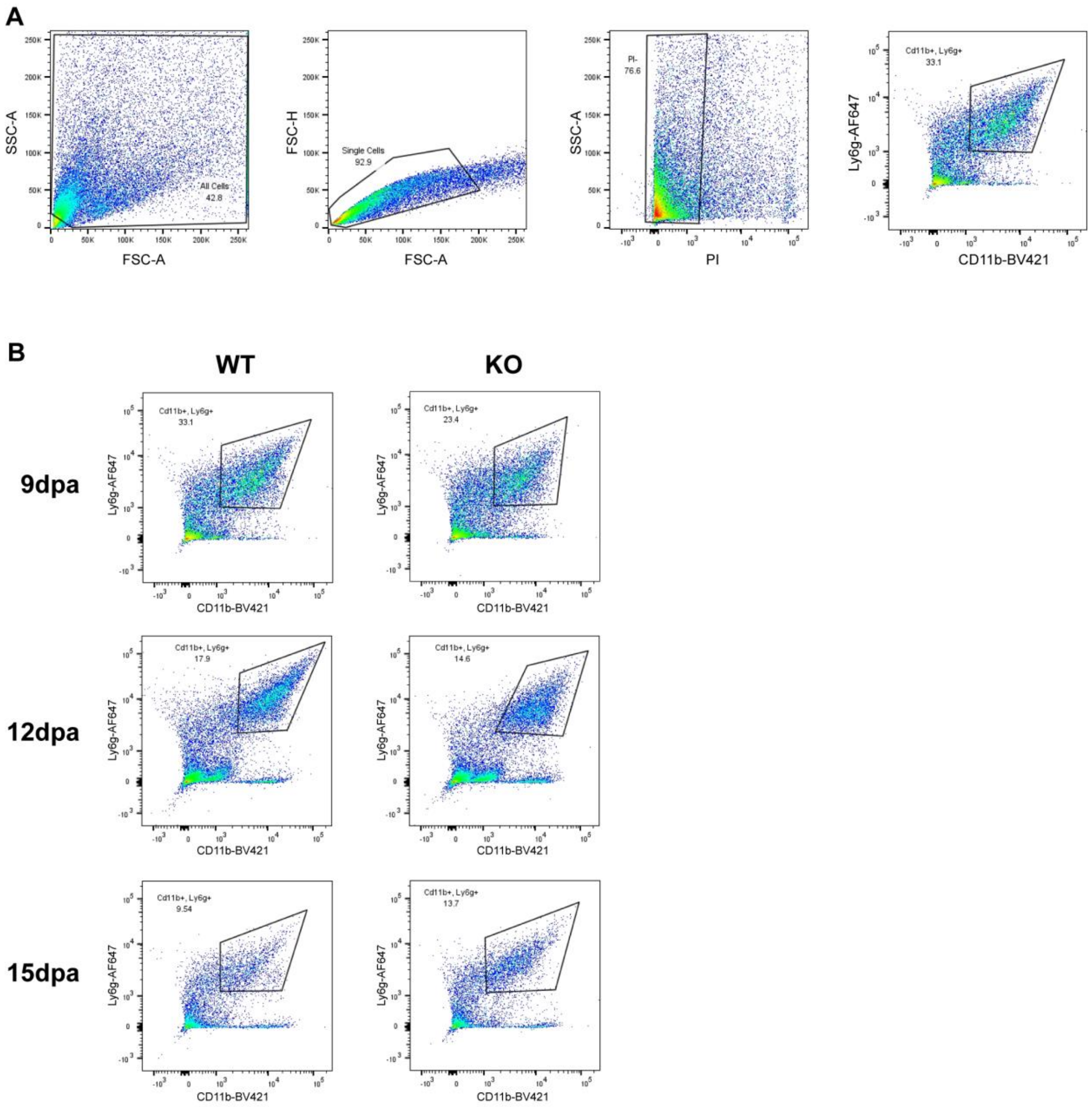
A
SSC-A
FSC-A
All Cells
42.8
FSC-H
Single Cells
92.9
PI-
76.6
PI
Ly6g-AF647
CD11b-BV421
Cd11b+, Ly6g+
33.1
B
WT
KO
9dpa
12dpa
15dpa
Cd11b+, Ly6g+
33.1
23.4
17.9
14.6
9.54
13.7

**Supplemental Figure 10**

**FACS analysis of neutrophils in wildtype and knockout tissue** (A) Sample gating strategy to quantify number of neutrophils. (B) CD11b-BV421 versus Ly6g-AF647 plots for WT and KO tissues at 9, 12, and 15dpa indicating the quantified region for $CD11b^{+}/Ly6g^{+}$ double-positive cells.

#### SUPPLEMENTAL TABLES

##### Supplemental Table 1. Primers

| | |
|---|---|
| **Genotyping primers:** | |
| Mest-F-Genotype | AATATGAGGATGTCTACACAACTCTTAA |
| Mest-R-Genotype | ACTCCAGCTTTAATTTGGCAGT |
| **qPCR primers:** | |
| GAPDH-F | AAGAGGGATGCTGCCCTTAC |
| GAPDH-R | CCATTTTGTCTACGGGACGA |
| Mest-Total-F | AAGCCATGTAAAAGCACAACTATCTC |
| Mest-Total-R | CCTACAAAGGCCTACGCATCTT |
| Mest211-F | GCGGCATGGGATAATGC |
| Mest211-R | CTACTTGGACCCACCACTCT |
| Mest210-F | GGGTAGAGAGAAAAAGTGTGGAA |
| Mest210-R | CCTCTAAGGAACAGCGACTTC |
| Mest202-F | CCCTGTGATCCGCAATCCT |
| Mest202-R | ACTACTGTCTGCATTTGGGCTATG |
| Perilipin-1-F | GGCCTGGACGACAAAACC |
| Perilipin-1-R | CAGGATGGGCTCCATGAC |
| PPARγ-F | CCAGAGCATGGTGCCTTCGCT |
| PPARγ-R | CAGCAACCATTGGGTCAGCTC |
| Col2a1-F | AGGGGTACCAGGTTCTCCATC |
| Col2a1-R | CTGCTCATCGCCGCGGTCCGA |
| Col10a1-F | ATGCCTTGTTCTCCTCTTACTGGA |
| Col10a1-R | CTTTCTGCTGCTAATGTTCTTGACC |
| Runx2-F | TCCACAAGGACAGAGTCAGATTAC |
| Runx2-R | TGGCTCAGATAGGAGGGGTA |
| **Bisulfite PCR primers:** | |
| CTCF-F | TGTTATTTTATATTTTTTGTAATAGGTGG |
| CTCF-R | CAAATAAAACCTATCCTACATACAAAAC |
| ExonA-F | GGTTAGGAGGTAATTTAAATTATTTGTT |
| ExonA-R | CACACATTATCTCATTTACTACTCAC |
| Exon1-5’-F | ATTTGGGGTTTAGGATTAGAGATTTATA |
| Exon1-5’-R | AAAAAAAACTACTACRCCTAAAAAAAA |
| Exon1-3’-F | AGTTGTTGTTTGTTGTTTTTGTTGT |
| Exon1-3’ -R | CCTTTAAATAAAAATTTTTACCTCCC |

**Supplemental Table 2. Differentially expressed genes upregulated in scRNAseq Mest-WT fibroblasts**

| gene | avg_log2FC | p_val_adj |
|---|---|---|
| Ccl2 | 2.34156423 | 8.31E-54 |
| Cxcl5 | 1.98077639 | 1.47E-36 |
| Cxcl1 | 1.95148461 | 7.87E-100 |
| Tnfaip3 | 1.91003983 | 1.29E-72 |
| Mest | 1.76395879 | 3.21E-75 |
| Cxcl2 | 1.59974693 | 2.10E-74 |
| Gm26771 | 1.47109463 | 1.00E-33 |
| Nr4a1 | 1.43910062 | 1.38E-40 |
| Nfkbia | 1.42959324 | 4.01E-121 |
| Zfp36 | 1.39327038 | 6.81E-68 |
| Dio2 | 1.38824246 | 1.85E-25 |
| Nfkb1 | 1.35876783 | 2.75E-34 |
| Ier3 | 1.32744243 | 5.24E-82 |
| Ptgs2 | 1.28631537 | 4.15E-21 |
| Ccl7 | 1.27844535 | 1.47E-20 |
| Ddx3y | 1.21368016 | 1.31E-80 |
| Pde10a | 1.17373904 | 1.45E-31 |
| Eif2s3y | 1.13980406 | 5.25E-69 |
| Slpi | 1.12664099 | 4.41E-39 |
| Serpina3n | 1.1173895 | 2.77E-28 |
| Mt2 | 1.10528856 | 3.52E-59 |
| Nfkbiz | 1.08716752 | 2.97E-77 |
| Sfrp2 | 1.07993783 | 2.01E-42 |
| Angptl4 | 1.01276328 | 8.40E-33 |
| Dusp1 | 0.99915841 | 2.17E-37 |
| Cebpd | 0.9990948 | 1.89E-20 |
| Maff | 0.99549165 | 1.77E-16 |
| Chl1 | 0.9738698 | 4.57E-19 |
| Ndrg1 | 0.95801991 | 2.53E-40 |
| Birc3 | 0.94343215 | 1.27E-14 |
| Pnrc1 | 0.93198878 | 1.22E-25 |
| Hspb1 | 0.93105983 | 9.10E-19 |
| Phlda1 | 0.92565355 | 1.61E-12 |
| Uty | 0.92034308 | 1.54E-55 |

| | | |
|---|---|---|
| Plpp3 | 0.91927962 | 2.57E-29 |
| Ddit4 | 0.91759515 | 2.18E-24 |
| Gem | 0.91206775 | 3.26E-48 |
| Nr4a2 | 0.89771788 | 1.58E-30 |
| Irf1 | 0.88708871 | 2.19E-31 |
| Rcan1 | 0.88254913 | 8.16E-28 |
| Pappa2 | 0.87704629 | 0.00202526 |
| Egr1 | 0.87572491 | 1.86E-54 |
| Arrdc3 | 0.87329321 | 2.94E-11 |
| Hspa1b | 0.87150305 | 9.72E-65 |
| Nfil3 | 0.86941835 | 1.47E-16 |
| Adamts1 | 0.86394997 | 1.19E-12 |
| Junb | 0.84822771 | 1.93E-71 |
| Cebpb | 0.84454004 | 4.25E-59 |
| Ntrk2 | 0.84403564 | 1.31E-18 |
| Cfb | 0.83207384 | 2.10E-13 |
| Fosb | 0.83118495 | 1.11E-16 |
| March3 | 0.82889961 | 6.70E-11 |
| Cadm2 | 0.78914276 | 0.000808 |
| Zfp36l1 | 0.78857663 | 3.78E-37 |
| Sdc4 | 0.78014257 | 1.16E-73 |
| Rspo4 | 0.77704207 | 4.02E-09 |
| Sik1 | 0.75977479 | 8.01E-16 |
| Hk2 | 0.7593632 | 1.10E-15 |
| Hspa1a | 0.75857196 | 8.13E-57 |
| Pim1 | 0.74465993 | 1.66E-16 |
| Klf9 | 0.74267027 | 9.79E-34 |
| Tnfaip2 | 0.73671805 | 2.37E-34 |
| Btg2 | 0.73374366 | 1.33E-27 |
| Hivep1 | 0.73006277 | 2.14E-14 |
| Tiparp | 0.72806166 | 7.90E-06 |
| Rbpms | 0.72473835 | 2.17E-26 |
| Schip1 | 0.72383564 | 0.00235524 |
| Ackr3 | 0.72129156 | 6.00E-11 |
| Ifrd1 | 0.71918814 | 4.16E-36 |
| Sod3 | 0.71809612 | 2.66E-97 |
| Slc6a17 | 0.71708275 | 1.17E-12 |
| Tgif1 | 0.71648015 | 3.31E-09 |
| Mt1 | 0.70387384 | 2.83E-53 |

| | | |
|---|---|---|
| Tnfsf11 | 0.70219356 | 2.31E-18 |
| Il11 | 0.69897883 | 9.51E-11 |
| Glis3 | 0.69808464 | 9.91E-66 |
| Cdkn1a | 0.69331898 | 1.46E-12 |
| Ier2 | 0.69249101 | 4.95E-32 |
| Igf2 | 0.68101283 | 0.00027552 |
| Rorb | 0.67999956 | 5.47E-18 |
| Sh3rf3 | 0.6793746 | 5.42E-07 |
| Jun | 0.67656001 | 1.15E-32 |
| Gadd45b | 0.67044861 | 4.25E-23 |
| Ets2 | 0.66916533 | 2.59E-16 |
| Dapk1 | 0.66844784 | 0.00345154 |
| Sox5 | 0.66612425 | 1.70E-07 |
| Igf2bp2 | 0.65610912 | 5.19E-09 |
| Tnfrsf11b | 0.65447818 | 0.00182813 |
| Lncpint | 0.64490747 | 3.77E-24 |
| Man1a | 0.64282503 | 5.10E-13 |
| Apoe | 0.64277737 | 0.00023825 |
| Steap4 | 0.64255647 | 5.33E-12 |
| Glul | 0.63932723 | 6.47E-06 |
| Hspa4l | 0.63529061 | 0.00053848 |
| Ptger4 | 0.63165589 | 2.56E-09 |
| Spp1 | 0.62805469 | 3.41E-15 |
| Bcl3 | 0.62612089 | 5.98E-13 |
| Btaf1 | 0.62409228 | 8.49E-50 |
| Ern1 | 0.62021302 | 6.69E-07 |
| Nfatc1 | 0.61885638 | 5.25E-13 |
| Mxd1 | 0.61751919 | 1.02E-09 |
| Fos | 0.6175066 | 4.18E-24 |
| Cxcl14 | 0.61740227 | 9.05E-21 |
| Cdh13 | 0.61597944 | 0.00268987 |
| Pfkfb3 | 0.61259018 | 7.12E-06 |
| Crim1 | 0.61176009 | 0.0009021 |
| Btg1 | 0.60919064 | 1.18E-49 |
| Pitpnc1 | 0.60839758 | 2.38E-22 |
| Slc16a1 | 0.6072917 | 6.93E-19 |
| Hipk2 | 0.60264817 | 1.05E-09 |
| Cmip | 0.59877163 | 1.43E-39 |
| Gfpt2 | 0.5983172 | 0.028104 |

| | | |
|---|---|---|
| Serping1 | 0.59502007 | 2.29E-30 |
| Slc4a4 | 0.5906213 | 8.29E-05 |
| Mgst1 | 0.58989248 | 8.29E-08 |
| St3gal1 | 0.58974132 | 1.05E-09 |
| Aff1 | 0.58905681 | 1.93E-26 |
| Pde1a | 0.58271423 | 5.00E-15 |

**Supplemental Table 3. Differentially expressed genes upregulated in scRNAseq Mest-KO fibroblasts**

| gene | avg_log2FC | p_val_adj |
|---|---|---|
| Matn3 | -1.0851626 | 2.82E-55 |
| Sept4 | -1.0217861 | 2.71E-43 |
| Npr3 | -0.9667224 | 1.05E-38 |
| Bcl11b | -0.9657123 | 3.66E-48 |
| Lsp1 | -0.903422 | 1.44E-76 |
| Sorbs2 | -0.8152891 | 2.06E-25 |
| Ecm2 | -0.7955849 | 5.36E-34 |
| Itm2a | -0.7941606 | 5.82E-64 |
| Lepr | -0.7495048 | 1.55E-19 |
| Omd | -0.7381966 | 1.42E-88 |
| Dlx6 | -0.73436 | 3.04E-29 |
| Col8a2 | -0.6880018 | 2.62E-62 |
| Ogn | -0.6796653 | 2.14E-47 |
| Cdh10 | -0.6789679 | 4.96E-33 |
| Hacd4 | -0.6767732 | 4.56E-48 |
| Fndc1 | -0.6711159 | 5.59E-74 |
| Pdlim2 | -0.6014694 | 3.78E-71 |
| Cdkn2b | -0.5976389 | 5.54E-18 |
| Gas2 | -0.5959082 | 1.56E-23 |
| Fat3 | -0.5907015 | 3.88E-13 |
| Dlx5 | -0.5868251 | 9.34E-31 |

**Supplemental Table 4. Differentially expressed genes for scRNAseq Mest-WT and KO integrated fibroblasts cluster-0**

| gene | avg_log2FC | p_val_adj |
|---|---|---|
| Cyp2f2 | 6.34080912 | 0 |
| Lcn2 | 5.65710861 | 0 |
| Dio3 | 5.43277833 | 0 |
| Nrg2 | 4.81359999 | 0 |
| Sfrp2 | 4.77934049 | 0 |
| Dio2 | 4.68736639 | 0 |
| Gm26771 | 4.63641059 | 0 |
| Apod | 4.60650856 | 0 |
| Cck | 4.56405487 | 3.53E-197 |
| Hhex | 4.55570143 | 0 |
| Pappa2 | 4.54854018 | 0 |
| Dio3os | 4.4685892 | 0 |
| Apoc1 | 4.46386155 | 0 |
| C4b | 4.40418619 | 0 |
| Steap4 | 4.39910113 | 0 |
| Tnxb | 4.37973759 | 0 |
| Slco2a1 | 4.33919485 | 0 |
| Lrg1 | 4.24875927 | 0 |
| Ccn3 | 4.2186837 | 0 |
| Vipr2 | 4.20443238 | 0 |
| Rspo4 | 4.19626175 | 0 |
| Rspo1 | 4.19050236 | 0 |
| Gpm6a | 4.16929537 | 0 |
| C3 | 4.07785118 | 0 |
| Corin | 4.07070636 | 0 |
| Gm26512 | 4.03487965 | 0 |
| Ramp2 | 4.02241096 | 0 |
| Serpina3n | 3.96859527 | 0 |
| Zcchc18 | 3.93960051 | 0 |
| Apoe | 3.92401807 | 0 |
| Cntn5 | 3.82567051 | 0 |
| Scara5 | 3.8242697 | 0 |
| Sorcs3 | 3.81151283 | 0 |
| Ntrk2 | 3.75273996 | 0 |

| | | |
|---|---|---|
| Cntn1 | 3.70973028 | 0 |
| Epha5 | 3.69113101 | 0 |
| Shisal1 | 3.68209453 | 9.99E-244 |
| Susd2 | 3.67330143 | 6.77E-209 |
| Trf | 3.66091695 | 5.52E-125 |
| Nrg3 | 3.65243197 | 0 |
| Cfb | 3.62162299 | 0 |
| Plpp3 | 3.51654181 | 0 |
| Mgst1 | 3.5162599 | 0 |
| Procr | 3.505636 | 0 |
| Emx2 | 3.50086002 | 1.67E-290 |
| a | 3.49031241 | 0 |
| Rarres2 | 3.45806947 | 0 |
| Prxl2a | 3.45626398 | 0 |
| Fbxo2 | 3.42935784 | 1.10E-240 |
| Ntrk3 | 3.42796389 | 0 |
| Hspb1 | 3.3561184 | 0 |
| Fos | 3.35322755 | 0 |
| Pcdh10 | 3.33578345 | 1.30E-238 |
| Cxcl5 | 3.30899532 | 3.16E-177 |
| Icam1 | 3.2882509 | 0 |
| Cebpd | 3.28086759 | 0 |
| Entpd2 | 3.24550033 | 0 |
| Efemp1 | 3.18688395 | 0 |
| Nr4a1 | 3.17747904 | 0 |
| Dusp1 | 3.16652227 | 0 |
| Wif1 | 3.15197592 | 0 |
| Ifitm1 | 3.14799256 | 0 |
| Syt13 | 3.08748364 | 0 |
| Prelp | 3.07716464 | 0 |
| Tst | 3.04654326 | 2.07E-260 |
| Maob | 3.04374672 | 0 |
| Calcrl | 3.04269749 | 9.95E-246 |
| Egfem1 | 3.03294006 | 0 |
| Csrnp1 | 3.01766371 | 2.86E-151 |
| Igf2 | 3.00591208 | 0 |
| Cybrd1 | 2.96877471 | 0 |
| Snhg11 | 2.96757921 | 7.71E-251 |
| Rgs6 | 2.9668774 | 0 |

| | | |
|---|---|---|
| Klhl32 | 2.96665456 | 2.32E-257 |
| Orm1 | 2.96071605 | 9.91E-146 |
| Ptger4 | 2.95014067 | 0 |
| Frmpd4 | 2.93166212 | 0 |
| Gstm1 | 2.91508403 | 0 |
| Isyna1 | 2.88718629 | 2.42E-157 |
| Hsd11b1 | 2.86704166 | 4.47E-106 |
| Gstt1 | 2.85016109 | 4.73E-293 |
| Ackr4 | 2.84966298 | 3.74E-234 |
| Rspo3 | 2.84436876 | 0 |
| Igfbp5 | 2.83323038 | 0 |
| Clu | 2.83145027 | 0 |
| Zfp36 | 2.8304276 | 2.68E-234 |
| Plxdc1 | 2.82987453 | 0 |
| Tnmd | 2.80726868 | 0 |
| Chl1 | 2.80516577 | 0 |
| Cxcl12 | 2.80393595 | 0 |
| Nog | 2.79260156 | 4.58E-194 |
| Sema3a | 2.77145576 | 0 |
| Dach1 | 2.75028485 | 0 |
| Gsta4 | 2.74646137 | 0 |
| Slc5a3 | 2.71154011 | 8.14E-185 |
| Spock1 | 2.68510964 | 0 |
| Cadm2 | 2.67729085 | 0 |
| Gch1 | 2.67095098 | 1.11E-108 |
| Gng8 | 2.66962851 | 1.36E-188 |
| Gna14 | 2.65349756 | 0 |
| Alx3 | 2.65092722 | 1.64E-165 |
| Emx2os | 2.64855272 | 1.19E-179 |
| Rorb | 2.64454768 | 0 |
| Dcn | 2.63732683 | 0 |
| Pcp4l1 | 2.63134256 | 2.50E-99 |
| 9330159F19Rik | 2.61004214 | 6.43E-196 |
| Gda | 2.60597569 | 3.79E-209 |
| Steap1 | 2.58779607 | 0 |
| Nr4a3 | 2.56091033 | 3.70E-177 |
| Draxin | 2.55761982 | 4.52E-186 |
| Dapk1 | 2.55565902 | 6.21E-218 |
| Kitl | 2.54485434 | 6.57E-145 |

| | | |
|---|---|---|
| Steap2 | 2.54229319 | 0 |
| Trpm3 | 2.53227928 | 0 |
| Stxbp6 | 2.51429478 | 0 |
| Wdr17 | 2.50825812 | 0 |
| Wfdc1 | 2.49653105 | 1.74E-271 |
| Ccl2 | 2.47518727 | 1.86E-41 |
| Cygb | 2.45899354 | 0 |
| Ccdc74a | 2.45835715 | 1.57E-152 |
| Fst | 2.44930957 | 0 |
| Ly6c1 | 2.44874445 | 0 |
| Pde10a | 2.44363192 | 1.53E-222 |
| Fosb | 2.43760147 | 1.20E-229 |
| Lbp | 2.42973937 | 0 |
| Hfe | 2.42618833 | 5.04E-289 |
| Ptger3 | 2.41743584 | 3.25E-203 |
| Cfap69 | 2.40663523 | 0 |
| Tox3 | 2.39982738 | 1.80E-198 |
| Hey2 | 2.3967081 | 5.06E-148 |
| C130021I20Rik | 2.39632451 | 7.40E-266 |
| Lmx1b | 2.39618546 | 0 |
| Add3 | 2.39583156 | 0 |
| Il34 | 2.37684923 | 2.14E-158 |
| Calml4 | 2.36807343 | 1.05E-102 |
| Btg2 | 2.36798278 | 0 |
| Cxadr | 2.3566763 | 0 |
| Nr4a2 | 2.3549928 | 5.67E-246 |
| Lamc3 | 2.34628445 | 2.62E-191 |
| Maff | 2.33461091 | 1.91E-145 |
| Cxcl14 | 2.33342659 | 0 |
| Serping1 | 2.33016048 | 0 |
| Ptgs2 | 2.32693391 | 1.80E-70 |
| Flrt3 | 2.28625505 | 3.75E-153 |
| Pnrc1 | 2.28607636 | 7.67E-297 |
| Tshz2 | 2.26386964 | 0 |
| Hspa1a | 2.25664503 | 0 |
| Junb | 2.24915588 | 0 |
| Ccdc3 | 2.24523285 | 4.12E-261 |
| Enpp2 | 2.2367081 | 0 |
| Rgs2 | 2.23470259 | 1.05E-95 |

| | | |
|---|---|---|
| Adh7 | 2.22335812 | 8.98E-89 |
| Sik1 | 2.21379477 | 1.24E-248 |
| S100a1 | 2.19123508 | 0 |
| Hspa1b | 2.18859523 | 0 |
| Gpm6b | 2.18858975 | 0 |
| Csmd3 | 2.18099325 | 8.25E-155 |
| Gm19522 | 2.18062862 | 1.80E-116 |
| C1s1 | 2.17499384 | 0 |
| Mapk13 | 2.16148544 | 1.34E-97 |
| Atf3 | 2.16070537 | 3.87E-41 |
| Gem | 2.16030782 | 2.78E-259 |
| Ly6a | 2.16005082 | 0 |
| Gfra1 | 2.15643466 | 3.85E-133 |
| Fgf2 | 2.143934 | 5.08E-214 |
| Olfml1 | 2.13336579 | 4.01E-159 |
| Nr3c2 | 2.11810255 | 4.24E-226 |
| Selenop | 2.10607419 | 1.41E-278 |
| Ednra | 2.10125124 | 0 |
| Ntng1 | 2.09338891 | 2.36E-216 |
| C1ra | 2.09225382 | 0 |
| B3galt1 | 2.09222241 | 1.05E-295 |
| Gm2a | 2.08687109 | 5.58E-194 |
| Ptprz1 | 2.08573721 | 2.35E-161 |
| Dpep1 | 2.07873678 | 7.82E-253 |
| Rab40b | 2.06990332 | 4.07E-132 |
| Masp1 | 2.06969016 | 0 |
| Sfrp1 | 2.05756797 | 1.01E-156 |
| Capn1 | 2.05145259 | 5.76E-128 |
| Tnfrsf11b | 2.05115969 | 1.14E-239 |
| Shox2 | 2.04874791 | 1.97E-138 |
| Txnip | 2.04526533 | 0 |
| Igfbp4 | 2.04345324 | 0 |
| Rom1 | 2.037574 | 1.90E-118 |
| Sybu | 2.02881173 | 6.25E-250 |
| Sgip1 | 2.02630488 | 0 |
| Scara3 | 2.01502406 | 0 |
| B3gnt5 | 2.00914722 | 5.09E-70 |
| Pear1 | 2.00337578 | 4.09E-115 |
| Adam33 | 2.00288684 | 2.72E-238 |

| | | |
|---|---|---|
| Grina | 1.99711815 | 1.07E-205 |
| Klf4 | 1.99566586 | 2.48E-125 |
| Nfkbia | 1.98570099 | 3.22E-219 |
| Cyb5a | 1.98139207 | 0 |
| Thsd4 | 1.98118084 | 5.28E-143 |
| Kctd1 | 1.97336123 | 0 |
| Cyp26b1 | 1.95984373 | 3.07E-185 |
| Trabd2b | 1.95021818 | 7.43E-288 |
| Sat1 | 1.94537858 | 0 |
| Glul | 1.94524232 | 5.42E-209 |
| Inpp4b | 1.94426141 | 8.35E-64 |
| Man1a | 1.94255827 | 0 |
| Gldn | 1.94022075 | 0 |
| Bambi | 1.93639993 | 6.94E-232 |
| Rnf180 | 1.9357463 | 2.17E-116 |
| Mllt3 | 1.92441982 | 0 |
| Daam2 | 1.92184498 | 1.92E-284 |
| Dnajc6 | 1.92162752 | 7.68E-68 |
| Meg3 | 1.91582216 | 1.35E-246 |
| Plscr4 | 1.91265185 | 8.47E-275 |
| Tmem204 | 1.91132635 | 3.89E-53 |
| Chd5 | 1.89676832 | 5.56E-95 |
| Cx3cl1 | 1.89576612 | 3.20E-120 |
| Cebpb | 1.88156623 | 0 |
| Cpne8 | 1.88127773 | 0 |
| Nos1ap | 1.87858333 | 5.46E-63 |
| Prr16 | 1.87852459 | 0 |
| Sned1 | 1.87572807 | 0 |
| Tnfaip3 | 1.86149707 | 7.99E-60 |
| Vdr | 1.8443144 | 2.70E-74 |
| Ndufa4l2 | 1.83960939 | 0 |
| Cxcl1 | 1.81758656 | 2.94E-37 |
| Rsph9 | 1.8127073 | 1.33E-52 |
| Nfil3 | 1.80706139 | 4.87E-78 |
| Slc6a6 | 1.80619749 | 0 |
| Hs3st1 | 1.80326163 | 1.16E-60 |
| Bcl2 | 1.80032218 | 2.40E-192 |
| Tmem132c | 1.79960991 | 1.66E-193 |
| Igf1 | 1.7989712 | 2.40E-164 |

| | | |
|---|---|---|
| Tmod2 | 1.79100423 | 2.35E-189 |
| Hs3st3a1 | 1.79067645 | 2.03E-90 |
| Grk5 | 1.78813942 | 2.67E-213 |
| Pcdh17 | 1.78352751 | 3.71E-122 |
| Ppp1r15a | 1.77537133 | 1.55E-165 |
| Flywch2 | 1.77091028 | 1.85E-86 |
| Id4 | 1.76282666 | 8.89E-48 |
| Pltp | 1.76224675 | 5.12E-193 |
| Ism1 | 1.76209621 | 1.44E-82 |
| Sall1 | 1.75599235 | 1.92E-78 |
| C630043F03Rik | 1.75479315 | 1.81E-95 |
| Fth1 | 1.75044212 | 0 |
| Col23a1 | 1.74649145 | 0 |
| Gadd45b | 1.74630964 | 5.92E-135 |
| Dab2 | 1.7406871 | 2.00E-285 |
| Acvrl1 | 1.73663013 | 1.14E-113 |
| Rab6b | 1.72925811 | 3.92E-55 |
| Birc3 | 1.72845567 | 1.28E-85 |
| Gm26532 | 1.72745051 | 4.12E-72 |
| Emc9 | 1.72417366 | 2.34E-50 |
| Hpse2 | 1.71431449 | 3.05E-266 |
| Klhl29 | 1.70691802 | 2.46E-110 |
| Twist1 | 1.70657878 | 0 |
| Fgfr3 | 1.69994541 | 3.64E-89 |
| Prxl2b | 1.69849308 | 1.52E-113 |
| Gm5089 | 1.6968869 | 3.80E-184 |
| Sp9 | 1.69056196 | 6.31E-65 |
| Pitx1 | 1.68858503 | 1.09E-196 |
| Pink1 | 1.68836255 | 8.79E-183 |
| Ier2 | 1.68258838 | 7.44E-165 |
| Vcan | 1.6805148 | 0 |
| Egr1 | 1.67996058 | 6.83E-117 |
| Ar | 1.67443267 | 9.38E-109 |
| Nkd2 | 1.67366396 | 9.71E-190 |
| Slc16a1 | 1.66913481 | 3.36E-152 |
| Inhbb | 1.65962727 | 8.87E-153 |
| Dpt | 1.65787365 | 4.05E-257 |
| Cd302 | 1.65568508 | 0 |
| Gfpt2 | 1.65126909 | 1.57E-133 |

| | | |
|---|---|---|
| Svep1 | 1.64996428 | 0 |
| Ahr | 1.64710209 | 5.31E-161 |
| Hsph1 | 1.64296406 | 4.91E-154 |
| Slc6a17 | 1.64009048 | 4.71E-143 |
| Sema3b | 1.63760548 | 2.29E-71 |
| Chst2 | 1.63445418 | 1.23E-86 |
| Adgra2 | 1.62792708 | 3.03E-283 |
| Zfp36l1 | 1.62315463 | 4.92E-198 |
| Aff2 | 1.62224723 | 2.29E-171 |
| Cdc14a | 1.621645 | 2.00E-98 |
| St6galnac5 | 1.61923786 | 2.12E-66 |
| Psen2 | 1.61496141 | 4.08E-83 |
| Ssbp2 | 1.61054341 | 1.10E-285 |
| Magi1 | 1.60929688 | 7.91E-150 |
| Rgs17 | 1.60810421 | 1.24E-59 |
| Sdc4 | 1.6068473 | 0 |
| Zic2 | 1.60675524 | 2.01E-107 |
| Prex2 | 1.60219917 | 6.00E-89 |
| Arrdc3 | 1.5993865 | 1.94E-82 |
| Dnajb1 | 1.59484737 | 3.37E-44 |
| 2200002D01Rik | 1.59421995 | 2.99E-49 |
| Pde4b | 1.59369154 | 3.75E-219 |
| Serpine2 | 1.59088314 | 0 |
| Ecscr | 1.58668973 | 1.05E-67 |
| Cdc42ep4 | 1.57894965 | 3.29E-70 |
| Abhd14b | 1.57803468 | 5.42E-73 |
| Bag3 | 1.57776877 | 2.13E-185 |
| Cox4i2 | 1.57758863 | 3.07E-86 |
| Il11ra1 | 1.57142648 | 0 |
| Mir99ahg | 1.57010482 | 0 |
| Igfbp6 | 1.56874983 | 1.34E-44 |
| Ptgis | 1.5680595 | 2.04E-301 |
| Galt | 1.56640575 | 1.81E-52 |
| Mxi1 | 1.55940603 | 1.42E-106 |
| S100a4 | 1.55622326 | 0 |
| Pde1a | 1.5553975 | 2.74E-208 |
| Ifi27 | 1.55424365 | 0 |
| 2310010J17Rik | 1.55421453 | 3.37E-86 |
| Atoh8 | 1.55027166 | 7.44E-37 |

| | | |
|---|---|---|
| Cdkn1c | 1.54714775 | 4.98E-89 |
| Epha7 | 1.54569947 | 8.70E-92 |
| Celf2 | 1.54549605 | 4.66E-91 |
| Tspyl4 | 1.54536161 | 4.66E-43 |
| Ndrg1 | 1.54218826 | 4.30E-200 |
| Slc16a2 | 1.54042355 | 0 |
| Ccl7 | 1.53591196 | 0.01798001 |
| Ifrd1 | 1.53276911 | 3.28E-226 |
| Bmp4 | 1.53055584 | 2.10E-211 |
| Crabp2 | 1.52888192 | 1.83E-70 |
| Efnb2 | 1.52774047 | 6.36E-222 |
| Tmem176a | 1.52721234 | 0 |
| 1500009L16Rik | 1.52620055 | 4.75E-38 |
| Irf1 | 1.5257205 | 5.94E-109 |
| Hebp1 | 1.52057668 | 4.55E-106 |
| Crlf1 | 1.51781108 | 0 |
| Msx1 | 1.51579043 | 0 |
| Igfbp2 | 1.5130004 | 3.09E-122 |
| Phyh | 1.51072471 | 8.83E-109 |
| Pde5a | 1.50940132 | 7.03E-99 |
| Reck | 1.50773565 | 4.68E-163 |
| Bmp2 | 1.50724807 | 1.53E-95 |
| Ddx3y | 1.50087661 | 7.17E-30 |
| Tmem158 | 1.49726815 | 3.84E-84 |
| Chp1 | 1.49638628 | 9.02E-205 |
| Olfm1 | 1.4949342 | 1.14E-171 |
| Zfp423 | 1.48863702 | 6.86E-74 |
| Lgals3 | 1.48632122 | 0 |
| Id2 | 1.48313185 | 6.94E-193 |
| Cd248 | 1.47370961 | 1.18E-246 |
| Pim3 | 1.46553429 | 7.08E-22 |
| Angptl4 | 1.46347146 | 9.41E-48 |
| Kctd8 | 1.45586449 | 5.37E-236 |
| Mtm1 | 1.45174718 | 2.27E-63 |
| Klf9 | 1.44777964 | 1.48E-141 |
| Mill2 | 1.44497284 | 3.66E-71 |
| Layn | 1.44453586 | 2.87E-110 |
| Nol3 | 1.44408476 | 1.11E-66 |
| Mvb12b | 1.44202353 | 3.95E-63 |

| | | |
|---|---|---|
| Zbtb4 | 1.44095412 | 5.10E-133 |
| Irak3 | 1.44055675 | 6.40E-42 |
| Npdc1 | 1.43844387 | 3.57E-158 |
| C1qtnf2 | 1.4370771 | 2.41E-62 |
| St3gal1 | 1.43450627 | 4.28E-103 |
| Cd81 | 1.43017447 | 0 |
| Jun | 1.42935511 | 3.09E-105 |
| Lefty1 | 1.42887347 | 3.57E-45 |
| Prkar1b | 1.42745338 | 2.59E-46 |
| Rerg | 1.41980777 | 4.40E-107 |
| Hsd17b4 | 1.41346813 | 9.26E-216 |
| Naaa | 1.41102862 | 4.33E-34 |
| Lmbrd1 | 1.41082822 | 2.77E-140 |
| Alx4 | 1.4084423 | 3.06E-166 |
| Nfkb1 | 1.39849891 | 3.10E-49 |
| Tfap2b | 1.39829195 | 3.52E-177 |
| Ccn1 | 1.39636353 | 9.84E-56 |
| Mcub | 1.39565005 | 3.63E-146 |
| Pnp | 1.39222615 | 4.37E-233 |
| Klf2 | 1.38858763 | 9.39E-11 |
| St6galnac3 | 1.38297695 | 3.84E-122 |
| Cd151 | 1.38192702 | 7.28E-125 |
| Ndrg2 | 1.3810225 | 4.28E-33 |
| Tmem176b | 1.3762319 | 0 |
| 9530026P05Rik | 1.36990262 | 1.39E-69 |
| Nfkbiz | 1.36683473 | 1.40E-71 |
| Gpc3 | 1.36560936 | 2.48E-67 |
| Zbtb20 | 1.36214526 | 0 |
| Ids | 1.35875347 | 1.26E-128 |
| Irs2 | 1.35155468 | 1.16E-37 |
| Gas6 | 1.34801259 | 2.47E-133 |
| Thy1 | 1.34466586 | 0 |
| Wnt11 | 1.33925066 | 7.43E-31 |
| Flot1 | 1.3321775 | 1.88E-167 |
| Lgals3bp | 1.33003767 | 3.43E-72 |
| Sema6d | 1.32698501 | 1.74E-66 |
| Pmp22 | 1.32489097 | 0 |
| Pgpep1 | 1.32457355 | 1.66E-23 |
| Ier3 | 1.32331911 | 1.54E-86 |

| | | |
|---|---|---|
| Ccnl1 | 1.32308141 | 4.92E-106 |
| Gja1 | 1.32065617 | 3.13E-231 |
| Smim1 | 1.3200819 | 4.31E-41 |
| Pbxip1 | 1.31558477 | 1.55E-178 |
| Vcam1 | 1.31468289 | 8.25E-220 |
| Gabbr1 | 1.31324329 | 3.35E-76 |
| Agtrap | 1.31316155 | 2.31E-35 |
| Fcgrt | 1.31301453 | 1.73E-238 |
| Fxyd1 | 1.3129196 | 3.62E-246 |
| Tcea3 | 1.31290432 | 3.80E-36 |
| Pitpnm2 | 1.31245587 | 1.53E-33 |
| Usp2 | 1.30902706 | 7.13E-32 |
| Abcb1a | 1.30615968 | 1.62E-25 |
| Mrps6 | 1.29882578 | 2.25E-74 |
| Lysmd2 | 1.29562339 | 8.28E-74 |
| Hes1 | 1.29254646 | 5.88E-97 |
| Adamts1 | 1.29248701 | 9.60E-35 |
| Tmem159 | 1.28998263 | 2.68E-109 |
| Acp2 | 1.2807492 | 5.63E-48 |
| Enc1 | 1.28037627 | 1.65E-35 |
| Scn7a | 1.27872945 | 3.69E-74 |
| Asap3 | 1.27835861 | 1.20E-63 |
| Crim1 | 1.27458111 | 4.22E-63 |
| Isg20 | 1.27360726 | 1.73E-26 |
| Dnaja4 | 1.27345767 | 3.11E-24 |
| Adi1 | 1.27196773 | 1.32E-29 |
| Ggt7 | 1.26950117 | 1.73E-44 |
| Zic5 | 1.26657976 | 2.81E-56 |
| Zfp703 | 1.25867561 | 1.72E-75 |
| Pdpn | 1.25781312 | 9.07E-147 |
| Sirpa | 1.25611298 | 1.03E-81 |
| Hspb8 | 1.25562057 | 8.87E-41 |
| Vegfa | 1.25350615 | 2.32E-89 |
| H2-D1 | 1.25248255 | 2.62E-196 |
| Cgnl1 | 1.25227752 | 7.73E-67 |
| Npr2 | 1.25087415 | 1.96E-60 |
| Sh3bp5 | 1.2498673 | 1.72E-64 |
| Slc22a17 | 1.24593566 | 6.99E-59 |
| Ophn1 | 1.24164442 | 3.49E-103 |

| | | |
|---|---|---|
| Ppp2r5b | 1.24053785 | 1.67E-23 |
| Tsc22d3 | 1.24005387 | 1.62E-28 |
| Vegfc | 1.23920727 | 5.02E-117 |
| Lum | 1.23778883 | 0 |
| Ror1 | 1.23606656 | 2.11E-141 |
| Lgals9 | 1.23570106 | 3.14E-114 |
| Nfia | 1.23494661 | 3.88E-195 |
| Rnase4 | 1.23484786 | 0 |
| Slc9a3r2 | 1.22971007 | 1.19E-26 |
| Rab4a | 1.22898886 | 2.85E-20 |
| Insyn1 | 1.22795599 | 4.58E-23 |
| Dync2li1 | 1.22750825 | 2.13E-38 |
| Btbd3 | 1.22744045 | 1.90E-36 |
| Csad | 1.2265231 | 3.29E-88 |
| Ptx3 | 1.22644165 | 9.53E-46 |
| Bmp7 | 1.22457094 | 3.42E-30 |
| Rnf220 | 1.2240137 | 2.52E-50 |
| Hs3st3b1 | 1.22390771 | 3.17E-38 |
| Prnp | 1.22379159 | 1.93E-298 |
| Boc | 1.22315475 | 2.25E-50 |
| Acyp1 | 1.22167512 | 1.08E-29 |
| Rab30 | 1.21968377 | 4.88E-69 |
| Mkx | 1.21593279 | 1.14E-121 |
| Ctsf | 1.21472435 | 1.39E-27 |
| Ltbp1 | 1.21138866 | 7.97E-214 |
| Pcsk5 | 1.21012352 | 1.72E-82 |
| Coq10b | 1.2087778 | 5.66E-85 |
| Bmyc | 1.20849662 | 8.43E-52 |
| Pcdh18 | 1.20844649 | 2.23E-73 |
| Mocs2 | 1.20748134 | 3.30E-216 |
| Jund | 1.20307432 | 8.33E-148 |
| Ccnd3 | 1.20152036 | 3.17E-144 |
| Apbb1ip | 1.19804198 | 5.73E-63 |
| Aldh3a2 | 1.19694049 | 4.01E-38 |
| Fhit | 1.19565781 | 1.77E-45 |
| Cacna1g | 1.19064465 | 1.81E-121 |
| Sertad1 | 1.18655539 | 3.47E-34 |
| Tgoln1 | 1.1857298 | 1.61E-155 |
| Tnfrsf1a | 1.18240804 | 9.31E-198 |

| | | |
|---|---|---|
| Id1 | 1.18126198 | 4.25E-67 |
| Sat2 | 1.17958516 | 1.10E-25 |
| Fbxo6 | 1.17429043 | 2.34E-19 |
| Stard5 | 1.17388283 | 2.02E-78 |
| Rnf167 | 1.17343458 | 3.51E-20 |
| Tcp11l2 | 1.17222943 | 1.39E-25 |
| Acvr1b | 1.16858549 | 2.52E-24 |
| Id3 | 1.16767127 | 2.06E-179 |
| Castor1 | 1.16605805 | 1.18E-33 |
| Hexim1 | 1.16604152 | 9.86E-60 |
| Tiparp | 1.15731349 | 1.19E-19 |
| Dnajb4 | 1.15627614 | 3.74E-76 |
| Scube1 | 1.15458547 | 2.57E-64 |
| Cib2 | 1.15397754 | 5.47E-45 |
| Pon2 | 1.15119065 | 2.94E-129 |
| Nkd1 | 1.14968566 | 1.48E-100 |
| Arrdc4 | 1.14765649 | 2.73E-55 |
| Mettl26 | 1.14374289 | 2.06E-12 |
| Dipk2a | 1.14294143 | 2.41E-108 |
| Ssc5d | 1.14275021 | 1.28E-37 |
| Pdgfra | 1.14023241 | 0 |
| Fosl2 | 1.13901527 | 1.32E-77 |
| Map1lc3a | 1.13428334 | 0 |
| Dcun1d3 | 1.13291952 | 1.32E-33 |
| Ltbp3 | 1.12956812 | 7.61E-131 |
| Dcun1d4 | 1.12851947 | 4.58E-09 |
| Lrp1 | 1.12828015 | 0 |
| Gaa | 1.12811381 | 5.92E-49 |
| Abca1 | 1.12045498 | 3.81E-76 |
| Igfbp7 | 1.11783896 | 0 |
| Ubc | 1.11260448 | 1.30E-214 |
| Gclc | 1.11176962 | 7.40E-28 |
| Dnajb9 | 1.1103562 | 1.38E-57 |
| Vamp2 | 1.10907155 | 5.87E-54 |
| Tmem205 | 1.10565261 | 4.68E-59 |
| Runx1t1 | 1.10518077 | 7.27E-147 |
| Ginm1 | 1.10311072 | 4.87E-143 |
| Tmem38b | 1.10130761 | 2.85E-15 |
| Scamp1 | 1.09841825 | 1.88E-62 |

| | | |
|---|---|---|
| Prps2 | 1.09838904 | 1.37E-19 |
| Wwtr1 | 1.09596703 | 6.03E-297 |
| Blvrb | 1.09204446 | 3.50E-143 |
| Emilin2 | 1.09063874 | 1.71E-66 |
| Smarca1 | 1.08644842 | 1.18E-22 |
| Cdkn1b | 1.08595445 | 5.42E-91 |
| Col8a1 | 1.08468477 | 1.05E-05 |
| Hgsnat | 1.08073426 | 8.66E-109 |
| Cpq | 1.07987096 | 7.70E-153 |
| Cachd1 | 1.07943896 | 9.71E-166 |
| Gm45338 | 1.07887586 | 8.57E-39 |
| Pgm2l1 | 1.0782045 | 9.00E-10 |
| Arhgap29 | 1.07700363 | 9.21E-39 |
| Ncoa7 | 1.07507022 | 3.74E-92 |
| S100a16 | 1.07484969 | 4.98E-181 |
| Dnajb2 | 1.07469076 | 9.19E-17 |
| Cmtm6 | 1.0735663 | 6.32E-28 |
| Il16 | 1.07289648 | 5.09E-39 |
| Fas | 1.07127376 | 3.70E-34 |
| Mr1 | 1.07101136 | 1.70E-13 |
| Stk40 | 1.06963562 | 2.26E-27 |
| Tmem63a | 1.06766866 | 7.97E-57 |
| Fuom | 1.0674642 | 2.15E-20 |
| Slc48a1 | 1.06623653 | 3.88E-71 |
| Gnptg | 1.0627977 | 1.10E-19 |
| Rsrp1 | 1.06254621 | 8.25E-234 |
| Tnfaip8 | 1.0625362 | 2.06E-63 |
| Tmem45a | 1.05955767 | 1.90E-122 |
| Plpbp | 1.05792431 | 2.02E-36 |
| Dhrs7 | 1.05695619 | 1.36E-66 |
| Arsk | 1.0566031 | 7.81E-14 |
| Cox6b2 | 1.05575274 | 1.17E-07 |
| Stat5a | 1.05502505 | 5.04E-23 |
| Hivep1 | 1.0546933 | 2.62E-10 |
| Mir22hg | 1.05467845 | 2.61E-53 |
| Erlin2 | 1.05320759 | 6.64E-13 |
| Pqlc3 | 1.0527826 | 1.29E-24 |
| Usp53 | 1.05277927 | 1.59E-12 |
| Gabarapl1 | 1.05219217 | 4.49E-19 |

| | | |
|---|---|---|
| Sod3 | 1.05036079 | 2.68E-185 |
| Aldh2 | 1.04974323 | 5.06E-155 |
| Ilvbl | 1.04966336 | 1.75E-12 |
| Pros1 | 1.04946643 | 7.76E-183 |
| Cdon | 1.04701632 | 2.75E-166 |
| Cbx4 | 1.04560092 | 2.46E-33 |
| Il1rap | 1.04498431 | 4.49E-32 |
| Ankrd12 | 1.04332774 | 2.34E-137 |
| Hapln1 | 1.04229937 | 1.78E-106 |
| Mxd1 | 1.04148878 | 1.23E-26 |
| Gstm5 | 1.04108021 | 1.12E-159 |
| S100a13 | 1.04062214 | 5.91E-217 |
| Crebrf | 1.03803853 | 2.11E-73 |
| Adgrl2 | 1.03708214 | 2.46E-157 |
| Lpar4 | 1.03694462 | 1.50E-14 |
| Thsd7a | 1.0357309 | 8.42E-112 |
| Akap7 | 1.03517874 | 2.56E-26 |
| Glis3 | 1.03317379 | 1.71E-169 |
| Bphl | 1.03213408 | 3.29E-20 |
| Uty | 1.03116822 | 7.86E-16 |
| Snx15 | 1.02484702 | 1.03E-28 |
| Ephx1 | 1.02350637 | 2.36E-15 |
| Atg2a | 1.02104606 | 2.56E-10 |
| Nsg1 | 1.01976021 | 7.02E-204 |
| Rhou | 1.01742842 | 5.81E-21 |
| Itgb8 | 1.01640647 | 9.37E-21 |
| Dcaf11 | 1.01633398 | 3.24E-25 |
| Zhx1 | 1.01403627 | 2.77E-31 |
| Tcn2 | 1.01120929 | 4.64E-53 |
| Plpp2 | 1.01109485 | 5.49E-29 |
| Paxx | 1.00969694 | 3.90E-27 |
| Etfrf1 | 1.00781633 | 5.76E-10 |
| Mt2 | 1.00703671 | 8.12E-10 |
| Cltb | 1.00509255 | 7.15E-121 |
